# Evaluating the applicability of kinship analyses for sedimentary ancient DNA datasets

**DOI:** 10.64898/2026.02.02.703194

**Authors:** Pnina Cohen, Sarah Johnson, Elena I. Zavala, Priya Moorjani, Viviane Slon

## Abstract

Kinship reconstruction in ancient populations provides key insights into past social organization and evolutionary history. Sedimentary ancient DNA (sedaDNA) enables access to deep-time human populations in the absence of skeletal remains. However, it is characterized by severe degradation and the potential mixture of genetic material from multiple individuals, raising questions about its suitability for kinship inference. Here, we use extensive simulations to evaluate the feasibility and limitations of kinship inference in sparse and damaged sedaDNA data, with a focus on Neandertals. We find that the main obstacle to accurate kinship inference in sedaDNA is the presence of multiple contributors to a given sample. To address this, we introduce a simple heterozygosity-based test to identify samples containing DNA from multiple individuals. Guided by these results, we analyze published Neandertal sedaDNA from the Galería de las Estatuas site to assess the practical limits of kinship inference in real sedimentary ancient DNA data. Together, our results define methodological considerations and practical limits for kinship inference in sedimentary ancient DNA.

## Introduction

The study of kinship in ancient human societies provides invaluable insights into the social structures, cultural practices, and evolutionary dynamics of early populations. Familial relationships have played a central role in shaping human history, underpinning genetic inheritance, social hierarchy, and the organization of communities. Previous research used skeletal remains and integrated burial context with genetic data to advance ancient kinship study. For instance, Haak *et al*. (2008)^1^, Fowler *et al*. (2022)^2^ and Rivollat *et al*. (2023)^3^ analysed ancient DNA (aDNA) from skeletal remains to infer familial relationships within Neolithic populations, revealing patrilineal kinship structures. Mittnik *et al*. (2019)^4^ examined burial sites from the Corded Ware Culture, concluding that complex family networks influenced burial arrangements and social status. Furthermore, Lalueza-Fox *et al*. (2010)^5^ and later Skov *et al*. (2022)^6^ showed that kinship studies - and hence also social dynamics - can also be inferred for ancient hominins such as the Neandertals.

Reconstructing kinship in ancient populations poses considerable challenges due to the fragmentary nature of the archaeological record, the degradation of biological material over time, and ancient and modern contaminants. These challenges are exacerbated at deep timescales (>50,000 years ago), particularly when only a few skeletal fragments may remain – if any. Recent work has demonstrated that ancient DNA can be obtained from sedimental samples (sedimentary DNA; sedaDNA) even when no skeletal remains are present^7–10^. This breakthrough offers an exciting new avenue for studying ancient human history, particularly at sites where other approaches relying on the preservation of human remains cannot be applied.

While sedaDNA expands the scope of kinship research, it also introduces new methodological difficulties. Compared to skeletal aDNA, human sedaDNA tends to be extremely low coverage, and frequently represents mixtures of DNA from multiple individuals. Under these conditions, kinship inference is constrained not only by sparse genetic information, but also by uncertainty arising from contributor mixture. As a result, understanding the limits of kinship inference in low-coverage, mixed ancient DNA data is essential for extending kinship analyses to sedaDNA contexts.

One approach to identify DNA mixtures has been to exploit variation in mitochondrial DNA within a single sample, enabling the detection of multiple maternal lineages even at very low coverage (e.g. Slon et al., 2017; Vernot et al., 2021; Vogel et al., 2024; Gelabert et al., 2025)^7,10–12^. In contrast, resolving mixtures using nuclear DNA typically requires either a small number of loci sequenced at very high depth and spanning long haplotypes (as in STR-based forensic analyses) or thousands of SNPs with near-complete genotyping and phasing^13,14^, conditions that are rarely met in sparse sedaDNA data.

In this study, we assess the performance of two kinship inference methods - KING and READ - and their limitations when applied to sedaDNA data and introduce a framework to detect mixtures of DNA profiles that constrain kinship inference in sparse ancient datasets. To do so, we simulated related and unrelated individuals from a Neandertal population. We then overlaid the simulated data with key features of ancient DNA, including reduced coverage, present-day human contamination, reference genome bias and substitutions resulting from deamination^15,16^. We additionally modeled sedaDNA-specific characteristics, including mixtures of multiple individuals within a sample and the presence of ancient faunal contamination. Building on these simulations, we developed an analytical pipeline for evaluating kinship inference from sedaDNA and applied it to a previously published dataset of Neandertal sedaDNA. This framework clarifies when kinship tools can be successfully applied for sedaDNA data, thus contributing to the broader field of aDNA research by validating new methodologies with real and simulated datasets.

## Methods

### Kinship Simulations

#### Creating a genotypes pool from a real Neandertal population

To simulate related individuals, we required a genotype pool of individuals from a single population, as differences between populations (e.g., allele frequency shifts, drift, or structure) would otherwise lead to spurious heterogeneity. We therefore used the genome-wide data of 12 Neandertal individuals that were excavated in Chagyrskaya cave in Siberia^6^, together with the high-coverage genome (∼28X) of the Chagyrskaya 08^17^ individual, all of whom were previously inferred the be from the same population^6^. Bam files were downloaded from http://cdna.eva.mpg.de/neandertal/ChagyrskayaOkladnikov/BAMS/Individuals/. After filtering for reads with mapping quality of at least 30, we ran *Glimpse2* to impute genotypes for each individual, using the 1000 Genome Project Phase III as a reference panel^18^ and the human genetic map GRCh37^19^ (see Supplementary Materials, Chapters 1 and 2, for an evaluation of imputation parameter choices for Neandertal genomes under low-coverage conditions). This resulted in a total of 78,152,284 genotyped SNPs. The imputed loci were filtered to retain only those with an imputed genotype probability (GP) of at least 80%, as was recommended by previous studies^20,21^, leaving 78,121,402 SNPs per individual. We then removed three individuals (*ChagyrskayaC, ChagyrskayaH* and *ChagyrskayaE*) of the four that were identified as possible first-degree relatives according to Skov *et al*. (2022)^6^, as first degree relatives in the genotypes pool could bias the simulations. We further filtered SNPs with missing data in any individual, retaining only variants with a MAF ≥ 5% to ensure that the minor allele would be present at least once in our dataset. This resulted in a final genotypes pool containing 931,274 SNPs, originating from ten individuals from Chagyrskaya cave.

#### Simulating a Neandertal family

*Sim1000G*^22^ is a genomic simulation framework that enables the simulation of individuals with an arbitrary (user-provided) pedigree structure using a catalogue of genotypes from a single population, under a recombination model in which crossover positions are drawn according to an empirical recombination map. We ran *Sim1000G* using the Chagyrskaya genotypes generated as detailed above, and the human genetic map for GRCh37^19^, to generate 14 “new” individuals from the Chagyrskaya population. Of these, 12 individuals constitute a three-generation family, and two have no genetic relatives (Figure 1). To construct genome-wide diploid genotypes with this pedigree structure, we used SNPs from all 22 autosomes, maintaining SNP counts approximately proportional to the chromosome lengths. Owing to the software constraints, we limited the number of SNPs per chromosome to a maximum of 36,000, resulting in a total of 513,204 SNPs sampled per simulated genome from the available genotype pool.

**Figure 1.**
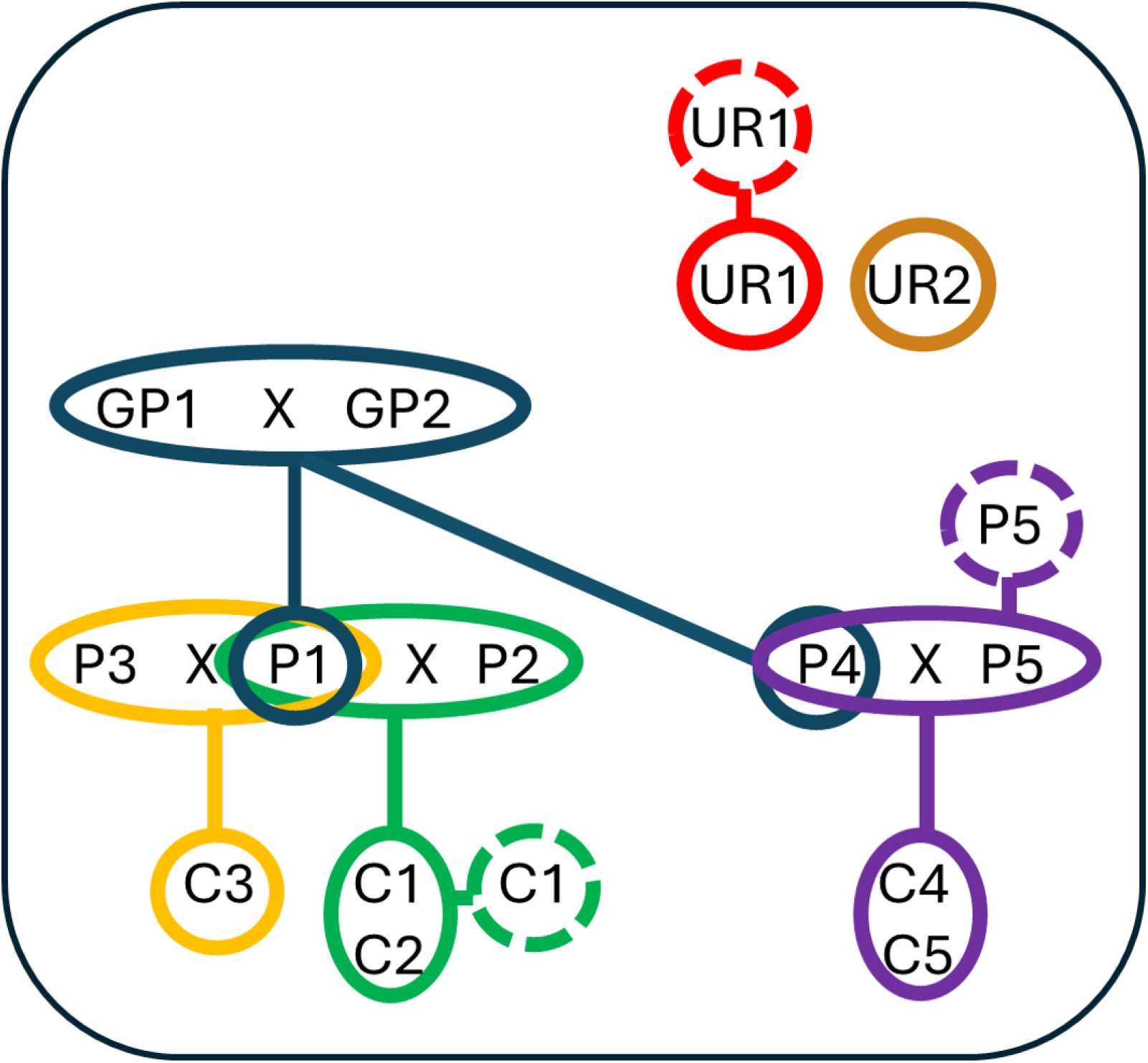
Simulated Neandertal genomes, comprising a three-generation family of 12 individuals and two additional unrelated individuals. Colors indicate sub-families based on first-degree relationships. Dashed circles represent duplicates of individuals with the same label. GP – grandparent; P – parent; C – child; UR – unrelated.

We independently simulated the individuals depicted in Figure 1 for a total of 100 replicates. For three individuals: a child (C1), a parent (P5), and an unrelated individual (UR1), we additionally generated exact copies. This was intended to model scenarios in which DNA fragments from the same individual are recovered across multiple samples, such as when distinct skeletal remains belong to a single person, or when DNA from the same individual is deposited in multiple sediment layers (due to leaching or sediment movement, for example). Altogether, the simulations encompassed individuals with kinship degrees ranging from 0 to 3, where 0 represents genetically identical individuals, 1 -first-degree relatives, 2 - second-degree relatives, and 3 for other relatives/unrelated individuals.

#### Simulating sedimentary ancient DNA data

Phased genotypes generated using *Sim1000G* were first ‘un-phased’ to better reflect available ancient DNA datasets. Specifically, we collapsed phased genotypes 0|1 and 1|0 to 0/1 and 0|0 and 1|1 to 0/0 and 1/1, respectively. To mimic key characteristics of ancient DNA, we introduced post-mortem DNA degradation patterns into the simulated genomes using the newly developed *ArchSim* simulator (GitHub repository: https://github.com/sarahaj32/archSim). *ArchS*im simulates the genotype patterns expected under several key processes relevant to ancient DNA data. Cytosine deamination is simulated by randomly converting homozygous reference calls at CpG transition sites to heterozygous at user-defined probabilities. Ancient faunal contamination is simulated by introducing ancestral alleles by randomly converting homozygous derived genotypes to heterozygous at a user-defined rate. Modern human contamination is simulated by randomly replacing endogenous genotypes with those drawn from a panel of modern human individuals. Reduced coverage and missingness were jointly simulated by sampling the sequencing depth from a negative binomial distribution at a user-specified mean depth and variance of 2, assigning this value to the DP parameter in the VCF file. Missing genotypes were then introduced when 0 reads were sampled. Heterozygous genotypes were converted to homozygous when 3 or fewer reads were sampled, with a bias towards selecting the reference allele of 0.55. Reference bias was further implemented by increasing the proportion of reference-matching alleles, up to 95% homozygous reference (0/0) calls. Additional implementation details for simulating features of ancient sediment DNA data are available in the *ArchSim* documentation.

In addition, studies of sedaDNA suggest that DNA recovered from a sedimentary sample can derive from more than one individual^7,8,10^. To assess the impact of mixture on kinship inference, we simulated two-person mixture pools by selecting different ratios of alleles from two individuals. For example, a 90:10 mixture would contribute nine alleles from one individual, and a single allele from the other at each locus. The genotype for each given locus was then established by randomly drawing two alleles from this pool.

The different sedaDNA-like features were applied to the simulated individuals independently across 100 replicates, either individually or in combination, to generate datasets of increasing complexity and degradation representative of sedimentary ancient DNA. Combined scenarios were generated to reflect the cumulative nature of degradation in real ancient DNA, with deamination, faunal contamination, modern human contamination, and missing data applied sequentially, such that each additional feature was introduced on top of the genotypes modified by the preceding step.

Specifically, we used *ArchSim* to implement (a) 5% or 10% cytosine deamination, (b) 5% or 10% ancient faunal contamination, (c) 5% or 10% modern human contamination, using five randomly-chosen present-day genomes as contaminants (three European and two East Asians from the 1000 Genomes Project Phase III^23^), (d) reduced sequencing coverage to 3X, 6X, 8X or 16X, and (e) 67-95% reference bias. Mixtures of individuals C1 and UR2 were generated in 90:10, 75:25 or 50:50 ratios.

### Kinship inference

We applied two kinship inference algorithms to all the simulated ancient genomes. The first was KING v2.2.4^24^, a popular kinship inference method which models the genetic distance between a pair of individuals as a function of their allele frequencies. The second was READ v2.0^25^, which was specifically developed for low coverage and degraded archaeological genetic data, requiring only pseudo-haploid genotypes as input. While many methods for kinship inference exist, we chose to focus on KING and READ because they are widely used, place minimal constraints on data type, and scale efficiently to large cohorts with modest computational requirements.

Following the application of the sedaDNA-like scenarios detailed above, the genotype inputs were quality filtered as follows, prior to being provided to the kinship inference programs. We only retained SNPs with MAF ≥ 0.05 and genotype calls in at least two individuals. Under missing data scenarios, we applied an additional minimum read-depth filter to reduce noise and allele dropouts associated with low coverage. For KING in particular, we required a minimum depth of 6 to minimize misclassification of diploid genotypes; while for READ, a minimum depth of 2 was applied. Consequently, the number of average SNPs shared between each sample-pair is much lower in KING compared to that of READ. To maintain comparability, we ran KING with a higher average coverage of 16X and 8X, which are approximately equivalent to the average coverage of 6X and 3X in READ, respectively. To reduce genetic linkage, no more than one SNP per 10,000 base-pair (bp) was allowed. We down sampled the SNPs dataset to 50,000, 10,000, 5,000, and 1,000 SNPs. For READ, pseudo-haploid genotypes were generated by randomly sampling one allele per locus.

KING was run with the --related option. For READ, the baseline pairwise mismatch rate between unrelated individuals (P0) was set to the median of all pairwise comparisons within the dataset, following the authors’ recommendation. Each simulation scenario was analyzed separately, such that all individuals within a dataset were generated under the same parameter set.

#### Assessing kinship inference

Pairwise relatedness between individuals was assessed using the kinship coefficient (K) estimated by KING and READ and interpreted according to the guidelines provided by each method for the first three kinship levels.

For KING, these are

- 0.5≥ K>0.354 for identical samples,
- 0.354≥K ≥0.177 for first degree relatives,
- 0.177≥K≥ 0.0884 for second degree relatives, and

For READ:

- 1≥K>0.375 for identical samples,
- 0.375≥K>0.1875 for first degree relatives,
- 0.1875≥K>0.09375 for second degree relatives.

Pairs with inferred K values below these thresholds were classified as unrelated. Although lower degrees of relatedness could, in principle, be inferred from such low values, we considered them too error-prone for reliable interpretation. Additionally, K values exceeding 0.5 for KING or 1 for READ were capped at 0.5 and 1, respectively, while negative K values were set to 0.

Inferred relationships were compared to the true underlying kinship in the simulated pedigrees to evaluate the accuracy of the inference algorithms, for varying data quality under the different scenarios simulated. For simulated mixtures, kinship was assessed separately relative to the known relationship with each of the two mixed individuals, C1 and UR2. In addition, we calculated the proportions of correctly inferred relationships, false negatives, and false positives. We consider a kinship inference to be accurate if it matches the simulated kinship between individuals; a false negative if the inferred kinship is more distant that the underlying simulation (e.g., inference of a first-degree relationship for two identical individuals); and a false positive if the inferred kinship is closer than the simulated scenario (e.g., inference of a first-degree relationship for a simulated second-degree one).

### Analysis of sedaDNA from Galería de las Estatuas site

#### Processing of bam files

We applied kinship analyses to published real data for sedaDNA from the Northern Spanish site of Galería de las Estatuas site described in Vernot *et al* (2021)^10^. We downloaded the unmapped bam files of the nuclear-DNA enriched sedimentary samples from EBI (https://www.ebi.ac.uk/ena/browser/view/PRJEB42656). 148 bam files were obtained for 105 sedimentary sub-samples collected from layers 2-5 in pit I and from layer 2 in pit II. These layers were found to contain ancient Neandertal DNA^10^. The bam files were processed using a standard approach for ancient sequences (Supplementary Chapter 5). Bam files belonging to the same sub-sample (identified by their library name) were merged before downstream processing, resulting in 105 samples (Supplementary Table S3).

#### Genotyping only putatively deaminated sequences

To ensure the use of authentic ancient DNA sequences, only forward sequences with C->T substitutions in the first three or the last three bases and reverse sequences with G->A substitutions in the first three or the last three bases were retained. These sites were then masked and not used in the downstream analysis. Each sample was genotyped individually using the *HaplotypeCaller, CombineGVCFs* and *GenotypeGVCFs* protocols with the default parameters of the GATK pipeline^26^ and underwent quality filtering (see details in Supplementary Chapter 5), resulting in the number of SNPs ranging between 0 and 282. The number of sequences or SNPs that remained after each processing stage are provided in Supplementary Table S3.

#### Mixture detection based on heterozygosity

To test whether observed heterozygosity exceeded that expected for a single individual, we applied a one-tailed binomial test. We defined *N* as the total number of sites carrying the alternative allele and genotyped as heterozygous (0/1) or homozygous alternative (1/1), and *S* as the number of heterozygous sites (0/1), such that *S/N* represents the observed heterozygosity. The baseline heterozygosity expectation for an individual in a given population was denoted as *p*_*0*_. The null hypothesis of no mixture was rejected when the probability of observing *S* heterozygous sites under a binomial distribution with parameters (*N, p*_*0*_) was significantly greater than expected for a single individual.

Since no reliable population-wide reference panel is available for Neandertals, we used three estimates of background heterozygosity: *p*_*0*_=0.21 derived from the simulations of heterozygosity using genotypes inferred for the Chagyrskaya population and shared by those inferred for the Estatuas population (median value heterozygosity for one individual); *p*_*0*_=0.36 and *p*_*0*_=0.4 derived from the simulations of heterozygosity using genotypes inferred for the Chagyrskaya population (median and maximal value respectively). More details in supplementary Chapters 4 and 5, supplementary Figures S7 and S8 and Supplementary Table S4.

#### Kinship analysis

Genotypes from all 105 Estatuas samples were filtered to retain SNPs shared by at least two samples with MAF ≥ 5%, a minDP = 2, and no more than one SNP per 10,000 bp. Samples with >80% missing data (n=65) were excluded, as were an additional 13 putative composite samples. These either showed evidence of multiple mtDNA contributors (Vernot *et al*. ^10^) or exhibited nuclear DNA heterozygosity inconsistent with a single individual under *p*_*0*_ = 0.4 in the binomial test described above (Supplementary Table S4). The final dataset comprised 27 samples with 8,895 SNPs, which were used for READ kinship analysis. Applying the stricter KING requirement of minDP = 6 yielded insufficient SNPs for reliable inference (26–393 shared SNPs per sample pair).

## Results

### Impact of sedaDNA characteristics on kinship inference

We simulated 14 individuals derived from the Chagyrskaya Neandertal population to evaluate kinship inference. Twelve of these simulated individuals were related (from identical individuals to second-degree relatives), and two were unrelated individuals from the same population (Figure 1). To model sedimentary ancient DNA characteristics, we applied multiple scenarios to these simulated genomes, including cytosine deamination, contamination from ancient faunal and present-day modern human sources, reduced coverage, and missing data. We ran READ^25^ and KING^24^ under single and combined parameter scenarios, and assessed their kinship inference success based on the inferred kinship coefficient (K) between each sample-pair compared against the true (simulated) underlying kinship. Results are shown in Figure 2. The corresponding proportions of correctly inferred kinship, of false negatives (i.e., inferred relatedness below the true level, indicating reduced sensitivity), and of false positives (i.e., cases inferred relatedness above the true level, indicating reduced specificity) are summarized in Supplementary Figure S3.

**Figure 2.**
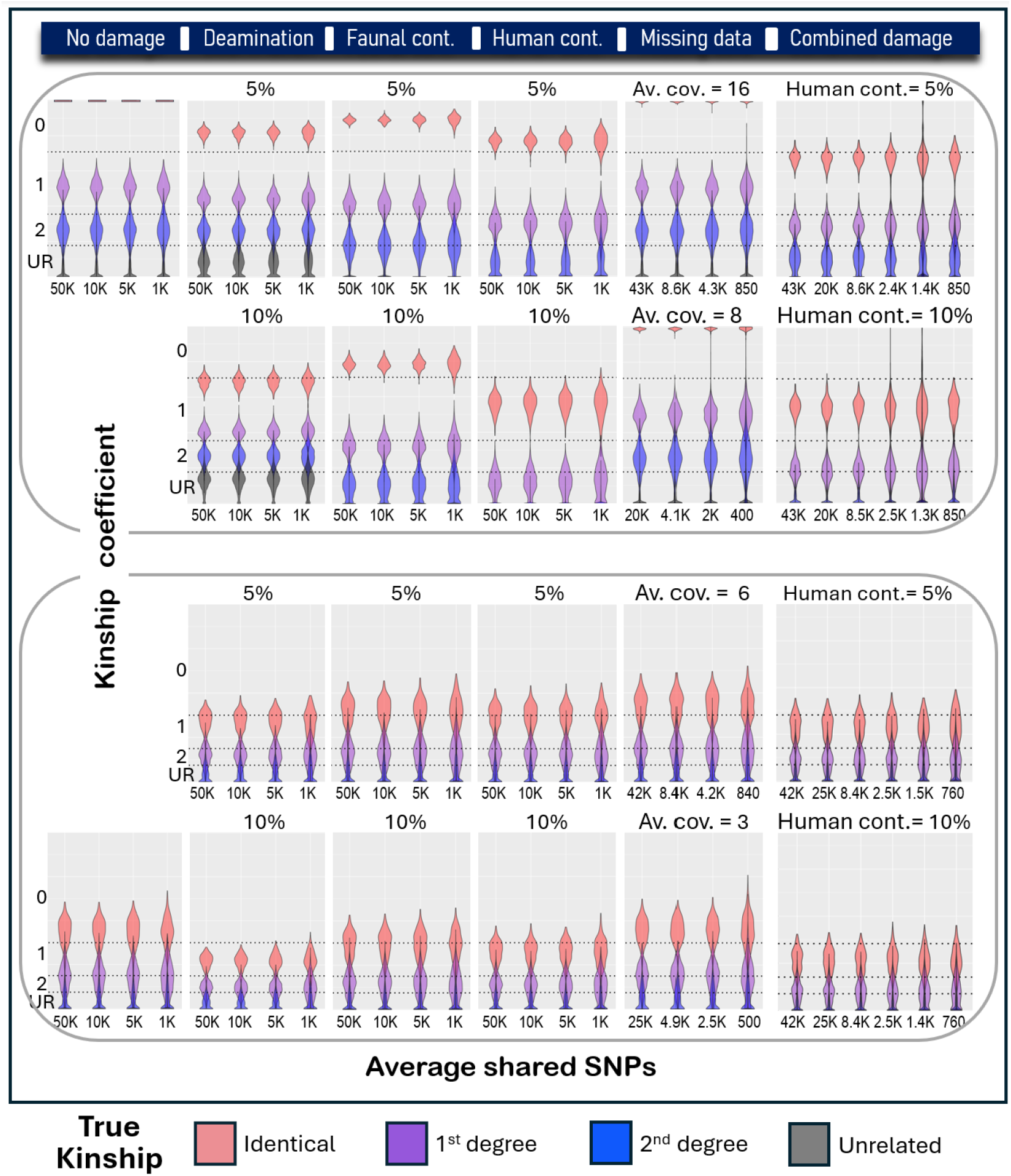
Kinship inference results from KING (top half) and READ (bottom half), obtained over 100 independently simulated families under various scenarios: deamination (5%/10%), ancient faunal contamination (5%,10%), modern human contamination (5%/10%), missing data (average coverages of 3X,6X/8X,16X), and the combination of the above. For the latter: deamination changes of 5% were simulated, followed by ancient faunal contamination of 5%, followed by modern human contamination of either 5% or 10%, followed by reduced coverage. Within each scenario, violin plots show the distribution of inferred kinship coefficient values across four bins of average shared SNP counts between individual pairs, rounded to the second significant digit (X-axis). Kinship classification thresholds (0: identical, 1: first-degree, 2: second-degree, UR: unrelated), as defined by each method’s guidelines, are marked on the Y-axis for reference. Plot colors indicate the true underlying degree of relatedness.

Under most scenarios, both KING and READ underestimated kinship as the severity of each simulated parameter increased, with false negatives exceeding false positives, indicating higher specificity than sensitivity. Using *KING*, 5% cytosine deamination had limited impact on kinship inferences, while 10% deamination increased both false positives where unrelated individuals were found related and false negatives in the inference for identical individuals. READ underestimated relatedness across all kinship classes at both simulated deamination levels. Faunal contamination showed a similar pattern, with KING remaining robust for identical individuals but increasingly underestimating closer relationships at higher contamination and READ exhibiting rising false-negative rates with increasing damage and decreasing relatedness. Both methods were robust to missing data, but with differing error profiles: at comparable numbers of shared SNPs, KING produced slightly more false positives but substantially fewer false negatives than READ. Achieving similar SNP counts, however, required higher initial coverage for KING due to its stricter filtering to avoid spurious homozygote calls, whereas READ’s pseudo-haploid framework is less affected by such issues.

We then evaluated the impact of mixtures of individuals from a single population being present in a single sample, by mixing DNA from child C1 and unrelated UR2 at varying proportions (Figure 3). This was performed both with and without introducing additional forms of sedaDNA characteristics, including modern human contamination of 5% (for 10% human contamination see Supplementary Figure S5). We then evaluated the inferred kinship of the mixed sample to all other individuals (excluding C1 and UR2) and compared these estimates to the true underlying kinship with either C1 or UR2. The proportions of accurate and misclassified inferences under these conditions are shown in Supplementary Figure S4.

**Figure 3.**
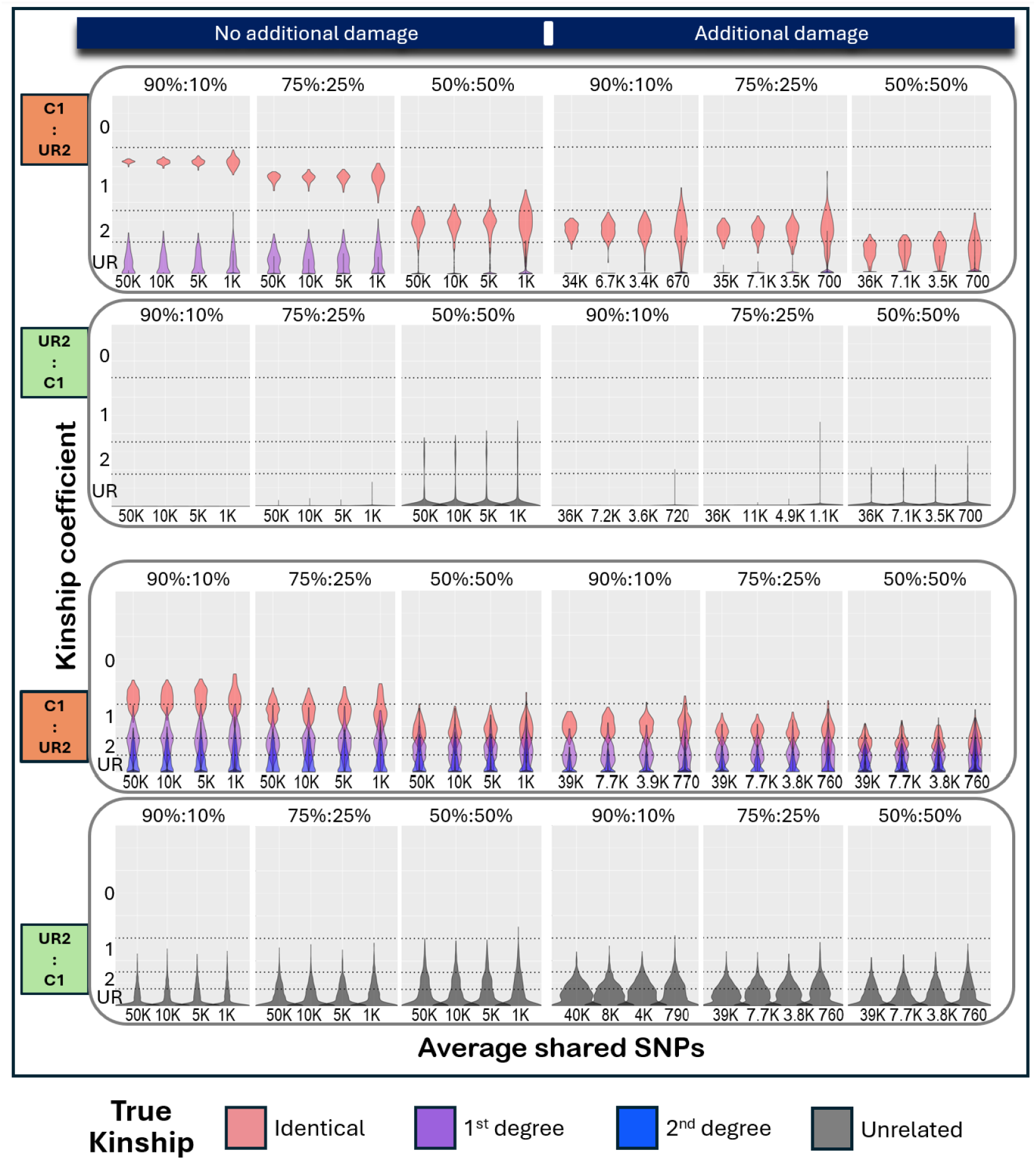
Kinship inference results from KING (top half) and READ (bottom half), obtained over 100 independently simulated families. Two individuals: C1 (child1) and UR2 (unrelated) were mixed in varying proportions to generate a composite sample. Violin plots display the distribution of the kinship coefficient values inferred only for pairs which include the mixed individual across various values of the average numbers of shared SNPs. **Left half**: No additional sedaDNA characteristics were simulated. **Right half**: Deamination changes of 5% were simulated, followed by ancient faunal contamination of 5%, modern human contamination of 5%, sample mixing and reduced coverage. **Odd rows (orange)**: Inferred kinship is compared with the underlying true kinship of C1. **Even rows (green)**: Inferred kinship is compared with the underlying true kinship of U2. The kinship classification thresholds (0: identical, 1: first-degree, 2: second-degree, UR: unrelated), as provided by each method’s guideline, are marked on the Y-axis as reference. Plots are colored by the true underlying degree of relatedness.

We find that mixture, either through modern human contamination or composite samples containing multiple individuals, had a pronounced impact on kinship inference. This effect was strongest for KING, while READ showed greater robustness, particularly for composite samples assessed from the related individual’s perspective (Child1). When evaluated from the unrelated individual (UR2), increasing mixture proportions led to elevated false positive kinship inferences (Figure 3; Supplementary Figures S4 and S5).

Finally, we evaluated how reference bias affects kinship inference (here, only READ method was applied), simulating up to 95% homozygous reference (0/0) calls. Due to filtering, reference allele concordance decreased to ∼70–85% of loci shared as 0/0 between sample pairs. We find that false positive rates increased under increasing reference bias, though in no case was the false positive rate above 20% (Supplementary Figure S6).

Overall, combining multiple types of sedaDNA characteristics resulted in substantial reduction in inference accuracy. The relative order of inferred kinship levels was generally preserved, albeit with increased overlap. Within each scenario, performance remained stable as the average number of shared SNPs per sample pair decreased from ∼50,000 to ∼5,000, even with accuracy was already compromised. A marked decline in performance was observed only when the number of shared SNPs dropped to ∼1,000, and in some cases to ∼500.

### A binomial test to identify signs of mixture of individuals in a single sample

Because sample mixture strongly impairs kinship inference (Figure 3; Supplementary Figures S4 and S45), we evaluated a heterozygosity-based test to detect samples containing DNA from multiple individuals. Simulations of mixed samples showed that heterozygosity levels could distinguish single-individual samples from mixtures in most tested scenarios (Supplementary Figures S7 and S8). This distinction was formalized using a binomial test comparing observed heterozygosity to that expected for a single individual. The binomial test with parameters (*N, p*_*0*_), where *p*_*0*_ is the expected heterozygosity in an individual and *N* is the number of evaluated sites, assesses whether the observed heterozygosity is significantly higher than expected under the null model. While this simple test distinguishes single-individual samples from mixtures, it does not provide further resolution as to the exact number of individuals contributing to a sample.

### sedaDNA of the Galería de las Estatuas site

To demonstrate mixed-sample detection and kinship reconstruction in real sedimentary DNA data, we analyzed 105 nuclear-DNA–enriched sediment samples from the Galería de las Estatuas^10^. To limit the impact of present-day human contamination, only putatively deaminated fragments (i.e., those with C→T substitutions at the first or last three bases) were retained. We then applied the heterozygosity binomial test to assess whether observed heterozygosity exceeded that expected for a single individual, using three background heterozygosity estimates derived from simulations (*p*_*0*_ = 0.21, 0.36, and 0.4; Supplementary Chapter 4).

We observed varied heterozygosity across samples, ranging from 0 to 0.63. Under the binomial test framework, 42 samples showed significantly elevated heterozygosity under *p*_*0*_ = 0.21 (p < 0.05), rejecting the single-individual null model, while eight samples remained significant under the strictest threshold (*p*_*0*_ = 0.4), consistent with composite samples. Vernot *et al*. identified nine samples with ≤10% modern human contamination based on mtDNA, six of which showed conflicting mtDNA positions indicative of multiple contributors. Of these, only one sample rejected the null hypothesis at *p*_*0*_ = 0.4, four did so only at *p*_*0*_ = 0.36 or 0.21, and one (A20287) did not reject the null hypothesis under any threshold (possibly due to the general low number of SNPs for this sample). Conversely, one sample which was consistent with a single contributor based on mtDNA, was identified by our test as a potential composite at *p*_*0*_ = 0.4 (A20281). This suggests that this test may identify mixed samples that cannot be identified by mtDNA differences alone. Full results are provided in Supplementary Table S4.

To enable kinship analysis with READ, we removed 65 low-coverage samples (>80% missing data) and 13 putative mixed samples (either identified by Vernot *et al*. or flagged by our binomial test at *p*_*0*_ = 0.4). After quality filtering, only eight of the 351 sample pairs shared more than 1,000 SNPs, and additional 15 pairs shared 800-1,000 SNPs. Among these, a single pair—library A16045 (Pit I, layer 3) and library A16112 (Pit II, layer 2)—was inferred by READ as containing DNA from second-degree relatives, based on 878 shared SNPs (Supplementary Table S5). These two samples were found to each likely originate from a single individual based on their mtDNA^10^. Based on our simulations, using READ at comparable levels of SNP information, unrelated individuals were misclassified as second-degree relatives in fewer than 15% of cases (Figure 2; Supplementary Figure S3). We further note that of these 878 shared SNPs, 610 (69.5%) were homozygous reference in both samples, a level not expected to produce false kinship inference rates above 10% (Supplementary Figure S6). Although these samples are derived from different stratigraphic layers dated to different time periods^27,28^, their mitochondrial DNA sequences were previously found to be very similar and their tip dates overlap^10^. Nevertheless, given the modest number of shared SNPs and the potential influence of contamination, reference bias, or other confounding factors, this inferred relationship should be treated with caution.

## Discussion

In this study, we developed a simulation framework to systematically evaluate the limits of kinship inference in highly degraded sedaDNA. By integrating real genetic data and modelling key sedaDNA features, including mixed samples, our approach provides a controlled setting to assess how individual and combined sedaDNA characteristics affect relatedness estimates. By identifying the conditions under which kinship inference remains reliable, this study clarifies the analytical potential of sedaDNA and broadens its use in reconstructing fine-scale social and demographic patterns in prehistory.

Among the typical sedaDNA characteristics simulated, deamination and ancient faunal contamination had minimal effect, particularly compared to the effects of mixtures of genetic information from distinct individuals – either the result of modern human contamination or in mixed samples containing DNA from multiple ancient individuals. Deamination and ancient faunal contamination primarily affect base calls at a subset of sites or introduce a small fraction of exogenous sequences, which tend to have only localized effects. In contrast, mixtures directly alter the genotype composition by introducing alleles from another individual, thereby distorting the identity-by-descent (IBD) patterns that kinship inference relies on.

Kinship accuracy by both KING and READ was generally underestimated in simulations with deamination, faunal and human contamination, and in composite samples when evaluated from the perspective of an individual related to others. As the severity of the sedaDNA-like characteristics increased, discrimination between second-degree relatives and unrelated pairs declined, subsequently reducing the differentiation between first-degree relatives and unrelated pairs. Identical pairs were inferred consistently and accurately, remaining reliable even in cases of severe degradation. Because false positives were rare and largely confined to unrelated and second-degree pairs, inference of first-degree relatives and especially identical individuals can generally be considered reliable in practical applications.

Only three scenarios produced substantial false positive rates: (1) high levels of deamination, particularly affecting KING; (2) strong reference bias (>80% reference allele calls); and (3) mixed samples evaluated from the perspective of an unrelated individual. The latter highlights a key challenge in sedaDNA analyses and underscores the need to detect mixed samples, while the former two emphasize the importance of rigorous data filtering and quality control prior to kinship inference.

Under identical conditions, KING generally outperformed READ for degradation scenarios including deamination and faunal contamination, and both methods showed comparable performance at similar numbers of shared SNPs despite reduced coverage. In contrast, READ consistently outperformed KING in the presence of modern human contamination or in mixed samples. READ estimates relatedness by comparing allele sharing within non-overlapping genomic windows, reducing sensitivity to uneven coverage and genotype uncertainty. Its reliance on pseudo-haploid genotypes eliminates the need for the strict minimum read-depth filtering required by KING, which can substantially reduce the number of SNPs available for analysis. Overall, we find that KING is preferable for use with relatively high-quality ancient DNA with minimal modern contamination or mixture of individuals, while READ is better suited for low coverage or pseudo-haploid data, highly degraded or mixed contexts – as is expected in sedaDNA.

We applied our methodology to real sedaDNA data from the Galería de las Estatuas cave, restricting our inference to sample pairs that shared at least 800 SNPs, and identified a single pair sharing a potential second-degree relationship. However, because the samples shared less than 1,000 SNPs and additional undetected contamination cannot be excluded, this inference should be treated with caution. Future applications of this approach to larger sedimentary datasets may provide more robust test cases for reconstructing kinship structures in ancient populations.

To address a major challenge in sedaDNA of mixed samples, we developed a simple binomial test that assigns a p-value for rejecting the single-individual null model given an expected heterozygosity (*p*_*0*_). Because *p*_*0*_ is population-dependent and difficult to estimate for ancient groups, as is illustrated by the Neandertal sedaDNA samples from the Galería de las Estatuas site^10^, which likely derive from several distinct populations, we applied the test across multiple *p*_*0*_ values. Resulting classifications were not always consistent with those reported by Vernot *et al*. based on mtDNA variation and genetic sex identification. Differences likely arise from the limited number of informative nuclear SNPs in some samples, and/or from the limited resolution of mitochondrial DNA for distinguishing individuals with identical haplotypes in others. Yet we consider the test a useful addition to the methodological toolkit for interpreting ancient sedimentary DNA.

Together with our kinship simulations, these findings provide a framework for evaluating the feasibility of relatedness inference in ancient populations under the extreme conditions typical of sedimentary ancient DNA. When extending these results to ancient modern human DNA recovered from sediments, several key differences should be considered. In particular, the smaller effective population size and reduced genetic diversity of Neandertals result in higher linkage disequilibrium and fewer informative polymorphisms for kinship inference, increasing sensitivity to noise and sample mixture. By contrast, ancient modern human populations generally exhibit higher genetic diversity and lower background relatedness, which are expected to improve the resolution of kinship inference under comparable data conditions. Consequently, our results from Neandertals likely represent a conservative lower bound for the performance that can be expected when inferring kinship from ancient modern human sedimentary DNA.

Patterns of relatedness are central to understanding human behavior, social organization, inheritance, and population dynamics. By demonstrating that mixture detection and kinship inference remain feasible even under highly constrained conditions, this study expands the scope of genetic analyses that can be applied to sedimentary DNA and other highly degraded ancient materials. Together, these results show that meaningful inferences about individual-level relationships can be recovered even in the absence of identifiable skeletal remains, opening new avenues for reconstructing social structures at sites previously considered genetically inaccessible.

## Acknowledgments

This study was funded by the Koret Berkeley-Tel Aviv University Initiative in Computational Biology and Bioinformatics (KBTI) (grant to P.M. and V.S.). V.S. acknowledges further support from the John Tempelton Foundation (grant #62571 to V.S.). E.I.Z. was funded by the Miller Institute in Basic Research Science at University of California, Berkeley and the Novo Nordisk Hallas-Møller Emerging Investigator Grant (NNF24OC0088862). The authors declare no conflict of interest.

Pnina Cohen would like to thank Laurent Briollais, Rasmus Henriksen, Yilei Huang and Harald Ringbauer, Benjamin Peter, Günther Torsten and Maróti Zoltán for their personal correspondence with her and their support using their respective software (though not all of them were eventually used in this study): sim1000G, NGSNGS, ancIBD, KIN, READ and correctKIN.

## Data deposition statement

All scripts used to generate simulated data are freely available in the Zenodo repository at https://doi.org/10.5281/zenodo.18450585. The aDNA simulator *ArchSim* is available on github at https://github.com/sarahaj32/archSim.

