## Supplementary material for "Evaluating the applicability of kinship analyses for sedimentary ancient DNA datasets"

### Supplementary Materials

#### **Chapter 1.** Evaluating the feasibility of imputation of Neandertal genomes using four imputation methods

Rationale

The simulation of related individuals from a Neandertal population requires genome-wide genotype data from multiple individuals. To date, more than 20 Neanderthal genomes have been published, however, all except three are low coverage^1–3^. We therefore opted to use imputation-based approaches to obtain complete genotype data from low-coverage individuals, so that these data would be suitable for downstream simulations. Imputation has been shown to be effective for the analyses of ancient human DNA datasets^4^ and its applicability to degraded modern human DNA for forensics purposes has also been assessed^5^. However, Neandertals represent a distinct hominin lineage that diverged from modern humans several hundred thousand years ago. This raises open questions about the feasibility and accuracy of imputing Neandertal genomes, motivating our systematic assessment.

##### Downsampling a high-coverage Neandertal genome for testing

We downloaded the high coverage (~52X coverage) “Altai Neandertal” (or “Denisova 5”) genome from southern Siberia^6^ from <http://cdna.eva.mpg.de/neandertal> in bam and VCF formats. We filtered the bam file for reads with a mapping quality of at least 30. Using the autosomes only, we ran *samtools* v1.6^7^ to randomly down-sample the bam files to an average coverage of 1X, 0.1X and 0.01X.

##### Imputation methods tested

The down-sampled bam files were imputed with four different software tools: *LinkImputeR*^8^, *Beagle*^9,10^ (two steps), *Quilt*^11^ and *Glimpse2*^12,13^. *Beagle* is a widely used imputation method; the other three are used for low coverage datasets, with Glimpse specifically designed for imputation of ancient DNA. LinkImputeR is a reference-free tool, while imputation with the last three tools was carried out using the human reference genome, the 1000 Genome Project phase III haplotypes as the reference panel^14^ and the hapmap genetic map GRCh37^15^. For all methods, imputation was executed separately and in parallel for each of the autosomes to reduce computation time.

##### Genotyping

Two of the imputation methods we tested - *LinkImputeR* and *Beagle* - impute based on a VCF genotypes file. We therefore used all 22 currently-available nuclear Neandertal genomes^2,3,6,16,17^ to create a VCF file. The bam files of these genomes were downloaded from <http://cdna.eva.mpg.de/neandertal/>. The individuals were genotyped using the *HaplotypeCaller*, *CombineGVCFs* and *GenotypeGVCFs* protocols with the default parameters of the *GATK* pipeline^18^ for the 22 autosomes, only using reads with mapping quality of 30 or more. The *GATK* pipeline was ran a total of four times: once with the full bam files and three more times for each of the reduced average coverages of 1X, 0.1X and 0.01X, resulting in 90,717,758, 91,033,475, 91,454,371 and 91,514,395 SNPs, respectively.

Six individuals with over 99% missing data were removed, leaving 16 individuals in the dataset. The SNPs were further filtered for a minimal allele frequency (MAF) of 3% and for a minimal read depth of six, except for the individual whose imputation quality is being examined (here – the Altai Neandertal), leaving 52,198,904, 54,321,304, 54,757,980 and 54,871,344 SNPs for the original coverage and the reduced coverages of 1X, 0.1X and 0.01X, respectively. These were used for downstream imputation with *Beagle*.

The SNPs were filtered further so only those with less than 30% missing data were kept, resulting in 8,419,869, 8,468,207, 8,498,781 and 8,502,778 SNPs for the original coverage and the reduced coverages of 1X, 0.1X and 0.01X, respectively. These were used in the downstream imputation with *LinkImputeR*.

##### Running *LinkImputeR*

*LinkImputeR* V1.1.4 was run over the VCF genotypes file with default parameters, imputing missing genotypes (for all individuals, not just Altai) using the other individuals in the VCF file as reference. Once imputation was done, the imputed loci were filtered so that only those with an imputed genotype probability of at least 80% were kept.

##### Running *Beagle* in two steps

To maximize *Beagle* performance, it was run in two steps^19,20^. *Beagle* version 4.0 (compiled on 27Jan18) was run over the genotypes file with the parameters *gl=true* (using genotype likelihoods), *impute=false* and the human genetic map GRCh37^15^. Like in the *LinkImputeR* run, missing genotypes were imputed using the individuals in the VCF file as reference. Over the results, *Beagle* version 5.0^21^ (compiled on 22Jul22) was run with the options: *gp=true, ap=false impute=true, burnin=20, iterations=200* using the 1000 Genome Project Phase III as the reference panel^14^ and the human genetic map GRCh37^15^ to complete imputation. Once imputation was done, only bi-allelic SNPs were kept with the minimal imputed genotype probability of 80%.

##### Running *Glimpse2*

*Glimpse*2 was run with default parameters over the Neandertal bam files using the 1000 Genome Project as the reference panel^14^ and the human genetic map GRCh37^15^. Once imputation was done, only bi-allelic SNPs were kept, as were those with an imputed genotype probability of at least 80%.

##### Running *QUILT*

*Quilt* V1.0.5 was run with the options: *--nGen=100* and *--buffer=1000* over the Neandertal bam files using the 1000 Genome Project as the reference panel^14^ and the human genetic map GRCh37^15^. Once imputation was done, only bi-allelic SNPs were kept, as were those with an imputed genotype probability of at least 80%.

##### Comparing the four imputation methods

The resulting imputed genotypes from all four methods were compared to the downloaded full genome sequence reconstructed using the high-coverage VCF file. The accuracy of imputation was assessed by calculating the average mismatch (or error rate) between the imputed alleles and the alleles in the full VCF. These were stratified by minor allele frequency (MAF) ranging between 0 and 0.45. The MAF was based either on the human reference panel genotypes (*Glimpse2*, *Beagle* and *Quilt*), or on the imputed genotypes themselves (of the 16 Neandertals genotyped together as described above; *LinkImputeR*). For clarification, bin of MAF=0 includes all SNPs while bin of MAF=0.01 includes all SNPs where the minor allele frequency is at least 1%, the bin of MAF=0.5 includes all SNPs where the minor allele frequency is at least 5%, etc.

For the higher coverages, *LinkImputeR* performed the best in terms of imputation accuracy out of the four methods tested (see the number of SNPs used in the evaluation and the error frequencies of the imputation results in Supplementary Table S1). However, with very reduced coverages (0.1x, 0.01X), not many SNPs remained post filtering (between 15 and 2,581 and between 3 and 90 SNPs, respectively), especially within the higher MAF bins. Given the expected low yields of DNA in ancient sediment samples^1^, this software is thus less likely to be widely applicable in the context of most sedaDNA research.

Of the remaining three methods, *Quilt* and *Glimpse2* performed similarly, and better than the two steps *Beagle*. However, *Quilt* requires a large amount of memory, and the full coverage imputation failed to complete, even when available allocated memory was increased to 256GB.

Therefore, overall, we found that *Glimpse2* outperforms the other three imputation algorithms under the tested scenarios, yielding higher accuracy and lower memory requirement. This is consistent with the findings of prior studies carried out on ancient and on degraded DNA^4,5^. Thus, we opted to perform further analyses only using *Glimpse2*.

**Supplementary Table S1.** Comparison between four imputation methods, performed for the high-coverage Altai Neandertal genome – using all available coverage, and after randomly down-sampling to lower coverage (X1, X0.1, X0.01). For each of those, the outcome of the comparison between the imputed genotypes and the data in the vcf file from the full-coverage genome are parsed by MAF bins ranging from 0 to 0.45 (accounting for both the major and the minor allele). For each software, the number of SNPs remaining in the analysis (lociNr) and the averaged error rate (avg.err) are reported.

|  | | **linkImputeR** | | **Glimpse2** | | **Quilt** | | **Beagle** | |
| --- | --- | --- | --- | --- | --- | --- | --- | --- | --- |
| Cov. | MAF | lociNr | avg.err | lociNr | avg.err | lociNr | avg.err | lociNr | avg.err |
| full | 0 | 7,900,049 | 0.000 | 77,261,721 | 0.002 | N/A | N/A | 72,455,080 | 0.012 |
|  | 0.01 | 7,900,021 | 0.000 | 11,861,961 | 0.006 | N/A | N/A | 10,040,488 | 0.059 |
|  | 0.05 | 7,889,656 | 0.000 | 6,725,299 | 0.008 | N/A | N/A | 5,449,085 | 0.076 |
|  | 0.1 | 762,528 | 0.001 | 5,157,747 | 0.008 | N/A | N/A | 4,121,230 | 0.080 |
|  | 0.15 | 594,656 | 0.001 | 4,150,881 | 0.009 | N/A | N/A | 3,288,660 | 0.084 |
|  | 0.2 | 97,125 | 0.002 | 3,361,656 | 0.009 | N/A | N/A | 2,646,297 | 0.087 |
|  | 0.25 | 85,678 | 0.002 | 2,682,587 | 0.009 | N/A | N/A | 2,101,363 | 0.089 |
|  | 0.3 | 42,856 | 0.002 | 2,078,126 | 0.010 | N/A | N/A | 1,622,342 | 0.091 |
|  | 0.35 | 27,671 | 0.001 | 1,522,852 | 0.010 | N/A | N/A | 1,184,504 | 0.092 |
|  | 0.4 | 23,099 | 0.001 | 998,028 | 0.010 | N/A | N/A | 774,125 | 0.093 |
|  | 0.45 | 7,914 | 0.001 | 495,054 | 0.010 | N/A | N/A | 382,540 | 0.093 |
| X1 | 0 | 790,955 | 0.002 | 75,828,154 | 0.010 | 74,930,237 | 0.009 | 76,517,755 | 0.024 |
|  | 0.01 | 790,880 | 0.002 | 10,725,615 | 0.048 | 9,941,351 | 0.048 | 11,173,053 | 0.129 |
|  | 0.05 | 790,115 | 0.002 | 5,854,169 | 0.059 | 5,283,810 | 0.059 | 6,186,965 | 0.173 |
|  | 0.1 | 78,573 | 0.017 | 4,427,190 | 0.062 | 3,961,026 | 0.063 | 4,705,894 | 0.187 |
|  | 0.15 | 60,547 | 0.021 | 3,528,894 | 0.065 | 3,135,870 | 0.066 | 3,769,709 | 0.198 |
|  | 0.2 | 11,764 | 0.087 | 2,836,562 | 0.068 | 2,506,346 | 0.069 | 3,043,599 | 0.207 |
|  | 0.25 | 10,474 | 0.090 | 2,250,431 | 0.070 | 1,978,643 | 0.071 | 2,424,540 | 0.214 |
|  | 0.3 | 5,761 | 0.114 | 1,735,033 | 0.072 | 1,519,798 | 0.073 | 1,875,913 | 0.219 |
|  | 0.35 | 3,906 | 0.111 | 1,266,287 | 0.073 | 1,106,439 | 0.074 | 1,372,420 | 0.222 |
|  | 0.4 | 3,303 | 0.108 | 827,273 | 0.075 | 722,041 | 0.075 | 898,766 | 0.225 |
|  | 0.45 | 1,164 | 0.079 | 409,031 | 0.076 | 356,170 | 0.077 | 445,620 | 0.227 |
| X01 | 0 | 2,581 | 0.019 | 74,597,231 | 0.012 | 74,016,951 | 0.012 | 76,983,861 | 0.040 |
|  | 0.01 | 2,566 | 0.018 | 9,523,697 | 0.065 | 9,024,470 | 0.070 | 11,536,081 | 0.229 |
|  | 0.05 | 2,566 | 0.018 | 4,811,204 | 0.085 | 4,461,100 | 0.095 | 6,432,360 | 0.336 |
|  | 0.1 | 426 | 0.090 | 3,508,958 | 0.092 | 3,227,453 | 0.104 | 4,891,095 | 0.387 |
|  | 0.15 | 310 | 0.108 | 2,723,015 | 0.098 | 2,489,843 | 0.111 | 3,918,112 | 0.428 |
|  | 0.2 | 147 | 0.184 | 2,143,958 | 0.103 | 1,952,027 | 0.117 | 3,165,595 | 0.463 |
|  | 0.25 | 136 | 0.180 | 1,670,942 | 0.108 | 1,517,371 | 0.122 | 2,522,980 | 0.494 |
|  | 0.3 | 79 | 0.209 | 1,273,362 | 0.111 | 1,150,584 | 0.125 | 1,953,533 | 0.518 |
|  | 0.35 | 47 | 0.160 | 919,834 | 0.114 | 829,730 | 0.128 | 1,431,379 | 0.538 |
|  | 0.4 | 38 | 0.145 | 597,572 | 0.116 | 539,364 | 0.129 | 937,773 | 0.552 |
|  | 0.45 | 15 | 0.100 | 292,736 | 0.119 | 263,026 | 0.132 | 465,332 | 0.558 |
| X001 | 0 | 90 | 0.078 | 72,755,331 | 0.012 | 74,022,826 | 0.012 | 77,000,505 | 0.041 |
|  | 0.01 | 89 | 0.079 | 7,524,949 | 0.071 | 9,029,819 | 0.071 | 11,550,852 | 0.238 |
|  | 0.05 | 89 | 0.079 | 2,870,788 | 0.093 | 4,462,542 | 0.096 | 6,443,457 | 0.353 |
|  | 0.1 | 39 | 0.141 | 1,736,004 | 0.099 | 3,228,123 | 0.104 | 4,898,019 | 0.409 |
|  | 0.15 | 32 | 0.156 | 1,148,753 | 0.107 | 2,490,837 | 0.111 | 3,922,537 | 0.457 |
|  | 0.2 | 25 | 0.200 | 788,174 | 0.118 | 1,954,015 | 0.117 | 3,168,499 | 0.500 |
|  | 0.25 | 19 | 0.237 | 546,443 | 0.129 | 1,519,281 | 0.123 | 2,525,571 | 0.536 |
|  | 0.3 | 16 | 0.250 | 375,986 | 0.137 | 1,154,118 | 0.127 | 1,956,281 | 0.566 |
|  | 0.35 | 10 | 0.250 | 253,022 | 0.145 | 832,145 | 0.130 | 1,434,225 | 0.589 |
|  | 0.4 | 6 | 0.083 | 156,936 | 0.150 | 540,786 | 0.132 | 939,910 | 0.606 |
|  | 0.45 | 3 | 0.167 | 74,702 | 0.157 | 264,410 | 0.135 | 466,597 | 0.615 |

#### **Chapter 2.** Further evaluation of *Glimpse2* and Neandertal imputation

##### Testing imputation accuracy on additional Neandertal genomes

Based on our preliminary results, suggesting *Glimpse2* outperforms the other three imputation algorithms, we performed subsequent tests using *Glimpse2* only. Two additional Neandertal genomes sequenced to high coverage: the Vindija 33.19 individual^17^ from Vindija Cave in Croatia (~30X coverage) and the Chagyrskaya 08 individual from Chagyrskaya Cave in Russia (~28X coverage)^16^ were downloaded from <http://cdna.eva.mpg.de/neandertal> in bam and VCF formats. These genomes were analyzed using the same processing and filtering steps as described for the Altai Neandertal genome, and imputed using *Glimpse2* with default parameters. The accuracy of imputation was then assessed as described above. Imputation accuracy is shown in Supplementary Figure 1, with detailed error and SNP metrics reported in Supplementary Table S2.

The accuracy of imputation was largely consistent across all three genomes within each down-sampling scheme. Unsurprisingly, in all three Neandertal genomes, imputation accuracy decreases at lower coverages, with the highest error rate observed at 0.01×, where errors reached up to ~18% at common variants (MAF >0.3). Error rates were also strongly influenced by allele frequency, with higher MAF associated with greater imputation error. Nevertheless, errors remained below 15% even at 0.01× coverage, provided MAF was below ~0.3. Overall, the results demonstrate the applicability of *Glimpse2* for imputing Neandertal genomes by leveraging the extensive human reference data available, if sequencing coverage is sufficiently high.

To our knowledge, this is the first attempt to impute Neandertal genotypes using a modern human reference panel, and our results show that this is a viable possibility when relatively sufficient genetic data is recovered. Notably, at 1× coverage, imputation of a Neandertal genome performs on par with ancient modern human datasets (error rates of ~5–7% for the former, compared to ~1.3–7.1% error for ancient humans outside Africa^4^).

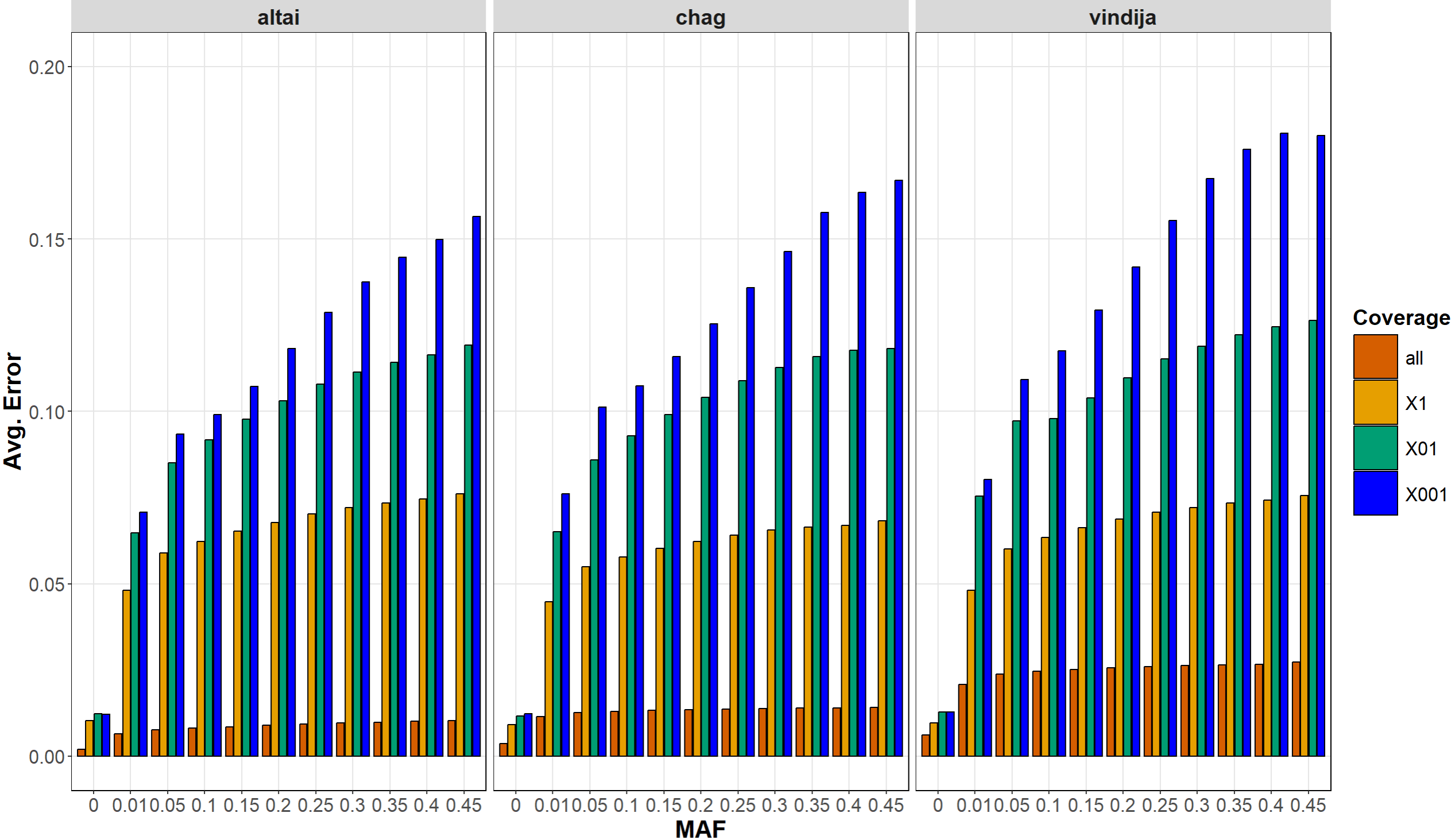
**Supplementary Figure 1**. Imputation results for three high-coverage Neandertal genomes using *Glimpse2*. For each Neandertal genome, the average imputation error is shown across different coverage levels: full dataset (no down-sampling), 1X, 0.1X, and 0.01X (randomly down-sampled). Errors are calculated by comparing imputed genotypes to the fully-sequenced genomes and are stratified into MAF bins ranging from 0 to 0.45 calculated over allele frequencies of the human reference genome. The error rate accounts for both the major and the minor allele.

**Supplementary Table S2**. Imputation results of *Glimpse2* for three high-coverage Neandertal genomes, either using the full dataset or after randomly downsampling to a reduced coverage (1X, 0.1X and 0.01X). For each coverage level and each Neandertal individual, the outcome of the comparison between the imputed genotypes and the data in the VCF file from the full-coverage genome are partitioned by MAF bins ranging from 0 to 0.45. To evaluate the performance of the software, the number of SNPs remaining in the analysis (lociNr) and the averaged error rates (avg.err) are reported.

| Individual | Coverage | MAF | lociNr | avg.err |
| --- | --- | --- | --- | --- |
| Altai | all | 0 | 77,261,721 | 0.002 |
|  |  | 0.01 | 11,861,961 | 0.006 |
|  |  | 0.05 | 6,725,299 | 0.008 |
|  |  | 0.1 | 5,157,747 | 0.008 |
|  |  | 0.15 | 4,150,881 | 0.009 |
|  |  | 0.2 | 3,361,656 | 0.009 |
|  |  | 0.25 | 2,682,587 | 0.009 |
|  |  | 0.3 | 2,078,126 | 0.010 |
|  |  | 0.35 | 1,522,852 | 0.010 |
|  |  | 0.4 | 998,028 | 0.010 |
|  |  | 0.45 | 495,054 | 0.010 |
|  | X1 | 0 | 75,828,154 | 0.010 |
|  |  | 0.01 | 10,725,615 | 0.048 |
|  |  | 0.05 | 5,854,169 | 0.059 |
|  |  | 0.1 | 4,427,190 | 0.062 |
|  |  | 0.15 | 3,528,894 | 0.065 |
|  |  | 0.2 | 2,836,562 | 0.068 |
|  |  | 0.25 | 2,250,431 | 0.070 |
|  |  | 0.3 | 1,735,033 | 0.072 |
|  |  | 0.35 | 1,266,287 | 0.073 |
|  |  | 0.4 | 827,273 | 0.075 |
|  |  | 0.45 | 409,031 | 0.076 |
|  | X01 | 0 | 74,597,231 | 0.012 |
|  |  | 0.01 | 9,523,697 | 0.065 |
|  |  | 0.05 | 4,811,204 | 0.085 |
|  |  | 0.1 | 3,508,958 | 0.092 |
|  |  | 0.15 | 2,723,015 | 0.098 |
|  |  | 0.2 | 2,143,958 | 0.103 |
|  |  | 0.25 | 1,670,942 | 0.108 |
|  |  | 0.3 | 1,273,362 | 0.111 |
|  |  | 0.35 | 919,834 | 0.114 |
|  |  | 0.4 | 597,572 | 0.116 |
|  |  | 0.45 | 292,736 | 0.119 |
|  | X001 | 0 | 72,755,331 | 0.012 |
|  |  | 0.01 | 7,524,949 | 0.071 |
|  |  | 0.05 | 2,870,788 | 0.093 |
|  |  | 0.1 | 1,736,004 | 0.099 |
|  |  | 0.15 | 1,148,753 | 0.107 |
|  |  | 0.2 | 788,174 | 0.118 |
|  |  | 0.25 | 546,443 | 0.129 |
|  |  | 0.3 | 375,986 | 0.137 |
|  |  | 0.35 | 253,022 | 0.145 |
|  |  | 0.4 | 156,936 | 0.150 |
|  |  | 0.45 | 74,702 | 0.157 |
| Chagyrskaya | all | 0 | 49,777,948 | 0.004 |
|  |  | 0.01 | 7,308,792 | 0.011 |
|  |  | 0.05 | 4,156,859 | 0.013 |
|  |  | 0.1 | 3,190,473 | 0.013 |
|  |  | 0.15 | 2,568,709 | 0.013 |
|  |  | 0.2 | 2,079,195 | 0.014 |
|  |  | 0.25 | 1,659,078 | 0.014 |
|  |  | 0.3 | 1,284,903 | 0.014 |
|  |  | 0.35 | 940,996 | 0.014 |
|  |  | 0.4 | 616,138 | 0.014 |
|  |  | 0.45 | 305,865 | 0.014 |
|  | X1 | 0 | 49,107,538 | 0.009 |
|  |  | 0.01 | 6,593,663 | 0.045 |
|  |  | 0.05 | 3,595,298 | 0.055 |
|  |  | 0.1 | 2,717,237 | 0.058 |
|  |  | 0.15 | 2,164,659 | 0.060 |
|  |  | 0.2 | 1,737,631 | 0.062 |
|  |  | 0.25 | 1,376,753 | 0.064 |
|  |  | 0.3 | 1,060,025 | 0.066 |
|  |  | 0.35 | 773,356 | 0.066 |
|  |  | 0.4 | 504,435 | 0.067 |
|  |  | 0.45 | 250,123 | 0.068 |
|  | X01 | 0 | 48,413,833 | 0.012 |
|  |  | 0.01 | 5,864,255 | 0.065 |
|  |  | 0.05 | 2,961,051 | 0.086 |
|  |  | 0.1 | 2,159,593 | 0.093 |
|  |  | 0.15 | 1,676,574 | 0.099 |
|  |  | 0.2 | 1,317,147 | 0.104 |
|  |  | 0.25 | 1,025,130 | 0.109 |
|  |  | 0.3 | 778,785 | 0.113 |
|  |  | 0.35 | 561,185 | 0.116 |
|  |  | 0.4 | 363,879 | 0.118 |
|  |  | 0.45 | 178,837 | 0.118 |
|  | X001 | 0 | 47,401,850 | 0.012 |
|  |  | 0.01 | 4,745,218 | 0.076 |
|  |  | 0.05 | 1,871,373 | 0.101 |
|  |  | 0.1 | 1,154,195 | 0.107 |
|  |  | 0.15 | 774,841 | 0.116 |
|  |  | 0.2 | 535,961 | 0.125 |
|  |  | 0.25 | 374,332 | 0.136 |
|  |  | 0.3 | 261,493 | 0.146 |
|  |  | 0.35 | 176,798 | 0.158 |
|  |  | 0.4 | 109,740 | 0.163 |
|  |  | 0.45 | 51,803 | 0.167 |
| Vindija | all | 0 | 49,315,128 | 0.006 |
|  |  | 0.01 | 7,214,013 | 0.021 |
|  |  | 0.05 | 4,094,771 | 0.024 |
|  |  | 0.1 | 3,140,610 | 0.025 |
|  |  | 0.15 | 2,527,358 | 0.025 |
|  |  | 0.2 | 2,044,689 | 0.026 |
|  |  | 0.25 | 1,630,909 | 0.026 |
|  |  | 0.3 | 1,262,865 | 0.026 |
|  |  | 0.35 | 924,510 | 0.026 |
|  |  | 0.4 | 605,345 | 0.027 |
|  |  | 0.45 | 300,520 | 0.027 |
|  | X1 | 0 | 48,955,090 | 0.010 |
|  |  | 0.01 | 6,495,000 | 0.048 |
|  |  | 0.05 | 3,517,840 | 0.060 |
|  |  | 0.1 | 2,652,354 | 0.063 |
|  |  | 0.15 | 2,108,403 | 0.066 |
|  |  | 0.2 | 1,689,115 | 0.069 |
|  |  | 0.25 | 1,337,147 | 0.071 |
|  |  | 0.3 | 1,029,205 | 0.072 |
|  |  | 0.35 | 750,334 | 0.073 |
|  |  | 0.4 | 488,899 | 0.074 |
|  |  | 0.45 | 241,651 | 0.076 |
|  | X01 | 0 | 48,002,505 | 0.013 |
|  |  | 0.01 | 5,383,386 | 0.075 |
|  |  | 0.05 | 2,406,428 | 0.097 |
|  |  | 0.1 | 1,597,167 | 0.098 |
|  |  | 0.15 | 1,209,953 | 0.104 |
|  |  | 0.2 | 946,518 | 0.110 |
|  |  | 0.25 | 734,405 | 0.115 |
|  |  | 0.3 | 556,558 | 0.119 |
|  |  | 0.35 | 400,315 | 0.122 |
|  |  | 0.4 | 258,276 | 0.125 |
|  |  | 0.45 | 127,749 | 0.126 |
|  | X001 | 0 | 47,439,548 | 0.013 |
|  |  | 0.01 | 4,781,239 | 0.080 |
|  |  | 0.05 | 1,897,877 | 0.109 |
|  |  | 0.1 | 1,176,251 | 0.117 |
|  |  | 0.15 | 796,511 | 0.129 |
|  |  | 0.2 | 555,272 | 0.142 |
|  |  | 0.25 | 392,016 | 0.155 |
|  |  | 0.3 | 275,486 | 0.167 |
|  |  | 0.35 | 187,101 | 0.176 |
|  |  | 0.4 | 115,577 | 0.181 |
|  |  | 0.45 | 55,969 | 0.180 |

##### Testing *Glimpse2* with European and East Asian only reference panel

Non-African populations were shown to share 1-3% percent of their genomes with Neandertals^22–26^. Therefore, we decided to check if the imputation results can be improved by reducing the reference panel to include only European and East-Asian populations, 503 and 504 individuals respectively, out of the original 2504.

We compared between the two imputation results (using complete reference panel and the reference panel of Europeans and East Asians) for all three Neandertal genomes and for all down-sampling schemes mentioned above. The performance of the imputation was largely worse using the restricted reference panel. Supplementary Figure S2 describes the relative differences between the results of the two imputation tests in terms of changes in the error rate and in the number of loci imputed. We therefore performed all additional analyses using the imputation scheme with *Glimpse2* and the full 1000 Genomes reference panel.

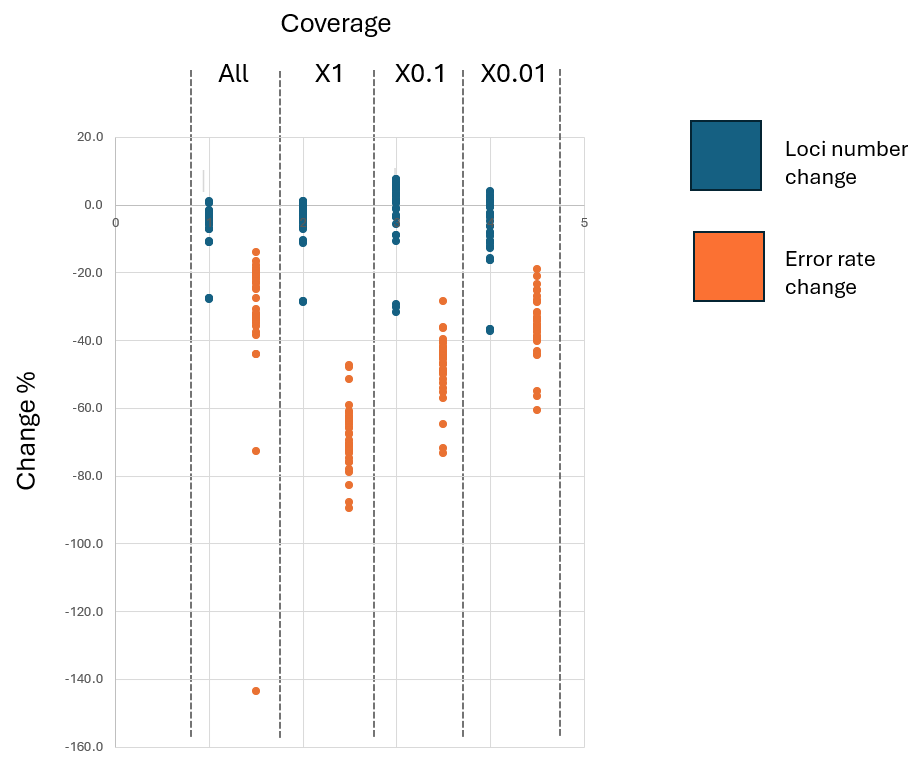
**Supplementary Figure S2.** Comparison of imputation performance in three fully sequenced Neandertals using the full 1000 Genomes reference panel (configuration I) versus only European and East Asian samples (configuration II). Differences in the number of imputed loci (blue) and average error rates (orange) are shown for configuration II relative to configuration I, with positive values indicating improved performance and negative values indicating reduced performance under configuration II.

#### **Chapter 3.** Kinship Simulations

##### Specificity and sensitivity analyses

In addition to the kinship inference results presented in Figures 2 and 3, we evaluated the accuracy of kinship inference in terms of sensitivity and specificity. Reduced sensitivity (false negatives) was defined as cases where kinship was missed altogether or inferred at a lower degree than the true relationship. Reduced specificity (false positives) was defined as cases where kinship was inferred despite not existing, or when the relationship was inferred at a higher degree than the true kinship. Supplementary Figure S3 presents this accuracy assessment for the scenarios in Figure 2, and Supplementary Figure S4 presents the same for the scenarios in Figure 3. Supplementary Figure S5 presents the results for simulations ran on mixed samples, with 10% present-day human contamination.

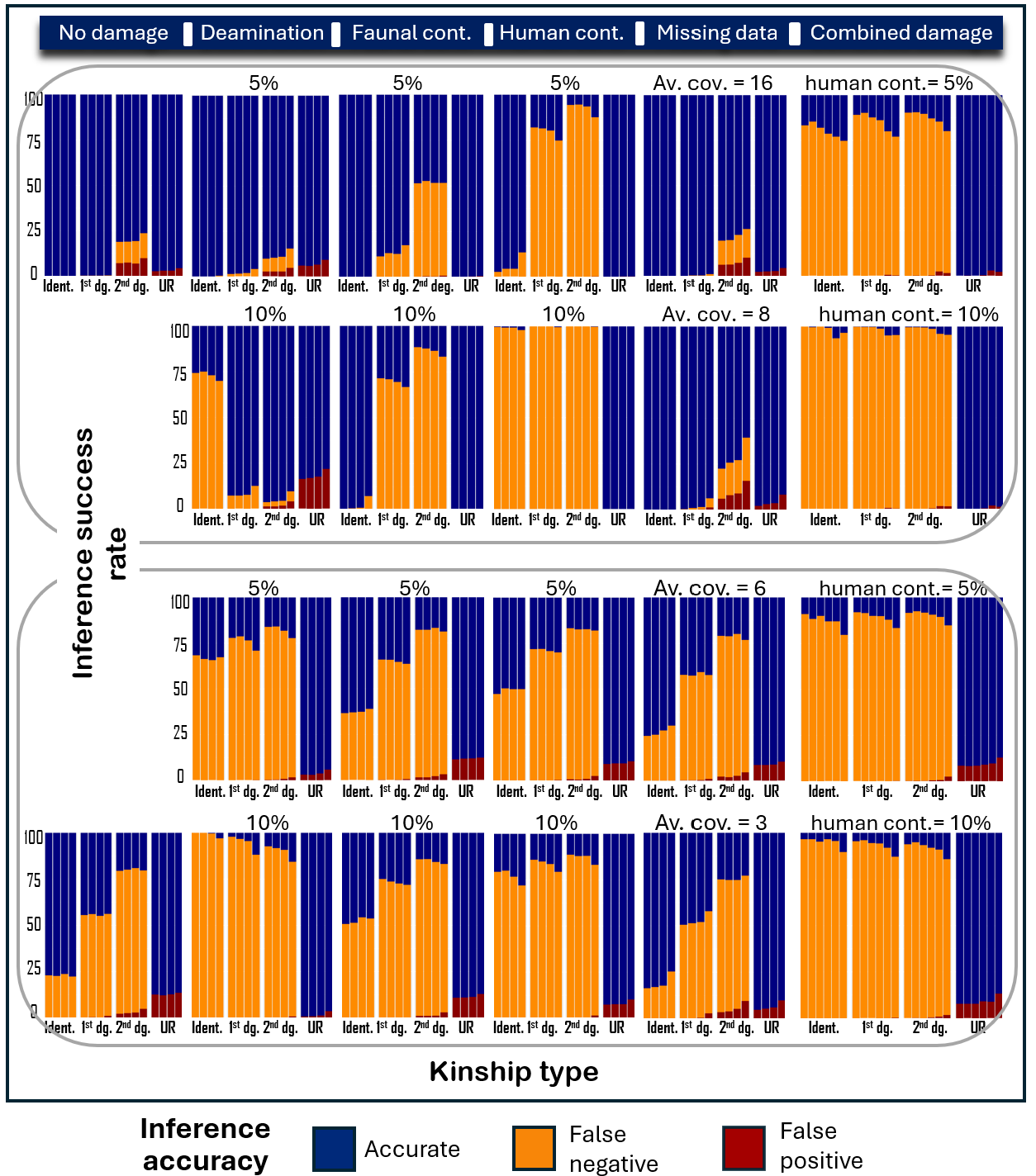
**Supplementary Figure S3**. Accuracy of kinship inference of *KING* (top half) and *READ* (bottom half), obtained by running each method over 100 independently simulated family under various scenarios: deamination (5%/10%), ancient faunal contamination (5%,10%), modern human contamination (5%/10%), missing data (average coverages of 3X,6X,8X,16X), and the combination of the above. For the latter: deamination changes of 5% were simulated, followed by ancient faunal contamination of 5%, followed by modern human contamination of either 5% or 10%, followed by reduced coverage. Within each scenario, four panels represent the four possible kinship types (identical, first-degree, second-degree and unrelated; X-axis), and within these, colored bars show the proportions of accurate, false negative and false positive inferences, across four values of the average SNPs number shared between individual pairs. The numbers themselves are not presented for visibility, but they can be found in main text Figure 2.

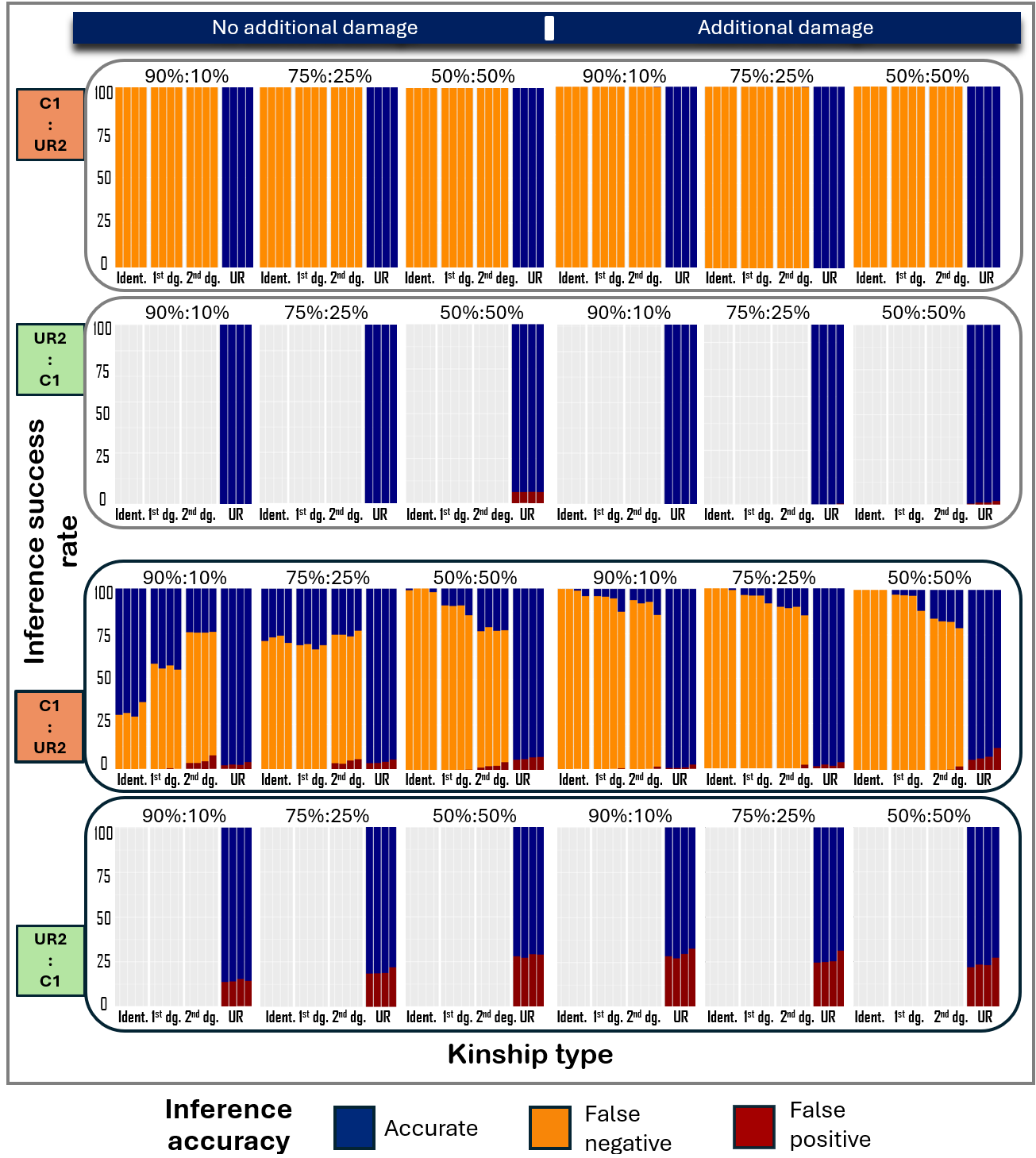
**Supplementary Figure S4.** Accuracy of kinship inference of *KING* (top half) and *READ* (bottom half), obtained over 100 independently simulated families. Two individuals: C1 (child1) and UR2 (unrelated) were mixed in varying proportions to generate a mixed sample. **Left half**: No additional sedaDNA characteristics were simulated. **Right half**: Deamination changes of 5% were followed by ancient faunal contamination of 5%, modern human contamination of 5%, sample mixing and reduced coverage. **Odd rows (orange)**: Inferred kinship is compared with the underlying true kinship of C1. **Even rows (green)**: Inferred kinship is compared with the underlying true kinship of U2. Four panels represent the four possible kinship types (identical, first-degree, second-degree and unrelated; X-axis). Within each panel, colored bars indicate the proportions of accurate, false negative and false positive inferences, across various values of the average SNPs number shared between individual pairs. The numbers themselves are not presented for visibility, but they can be found in main text Figure 3.

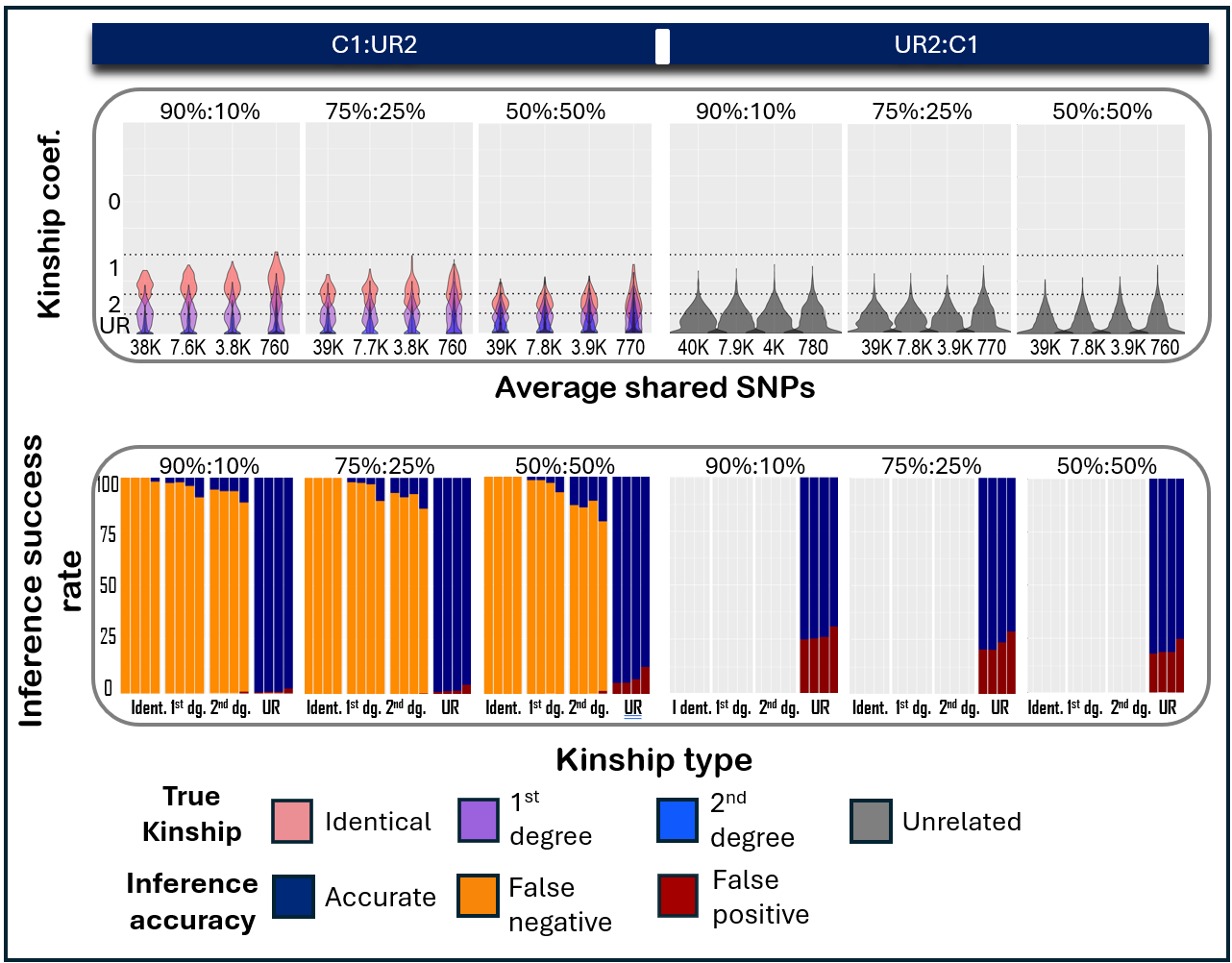
**Supplementary Figure S5.** Kinship coefficient and inference accuracy obtained by running *READ* over 100 independently simulated families. Two individuals: C1 (child1) and UR2 (unrelated) were mixed in varying proportions to generate a mixed sample. In addition, deamination changes of 5% were followed by ancient faunal contamination of 5%, modern human contamination of 10%, sample mixing and reduced coverage. **Left half**: the inferred kinship is compared with the underlying true kinship of C1. **Right half**: the inferred kinship is compared with the underlying true kinship of U2. **Top row**: Violin plots display the distribution of the kinship coefficient values inferred only for the composite sample across various values of the average numbers of SNPs shared between individual pairs. **Bottom row**: Accuracy of kinship inference. Four panels represent the four possible kinship types (identical, first-degree, second-degree and unrelated; X-axis). Colored bars indicate the proportions of accurate, false negative and false positive inferences, across various values of the average SNPs number shared between individual pairs.

##### Effect of reference bias on kinship inference

Monitoring the number of SNPs shared between sample pairs is clearly essential for assessing the reliable kinship inference, to the extent that both *KING* and *READ* explicitly report this value. For *READ*, a previous study estimated that at least ~1,000 shared SNPs are needed for dependable inference^27^, and our results suggest that reliable classification is possible even at somewhat lower thresholds. Nevertheless, reference bias introduces an additional and orthogonal source of error. When allele calls are preferentially inferred as the reference allele: due to extremely low coverage, imputation errors, or substantial genetic divergence between the study samples and the reference genome (e.g., Neandertals with respect to the human reference), the resulting genotypes may become artifactually homogenized. This artificially inflates apparent similarity between samples, leading to upward bias in inferred kinship coefficients.

To quantify this effect, we used *ArchSim* to introduce controlled levels of reference bias rates into our simulated genotypes, ranging between 70% and 95%, while ensuring that the minimum number of shared loci between sample pairs after filtering remained above 1,000 SNPs. We then assessed the impact of reference bias on kinship inference accuracy. As shown in Figure S6, reference bias has a clear and measurable effect on kinship accuracy, which is partly alleviated but not not fully mitigated by standard filtering procedures. We therefore conclude that reference bias should be explicitly assessed and accounted for when interpreting kinship estimates from low-coverage or evolutionarily divergent genomes.

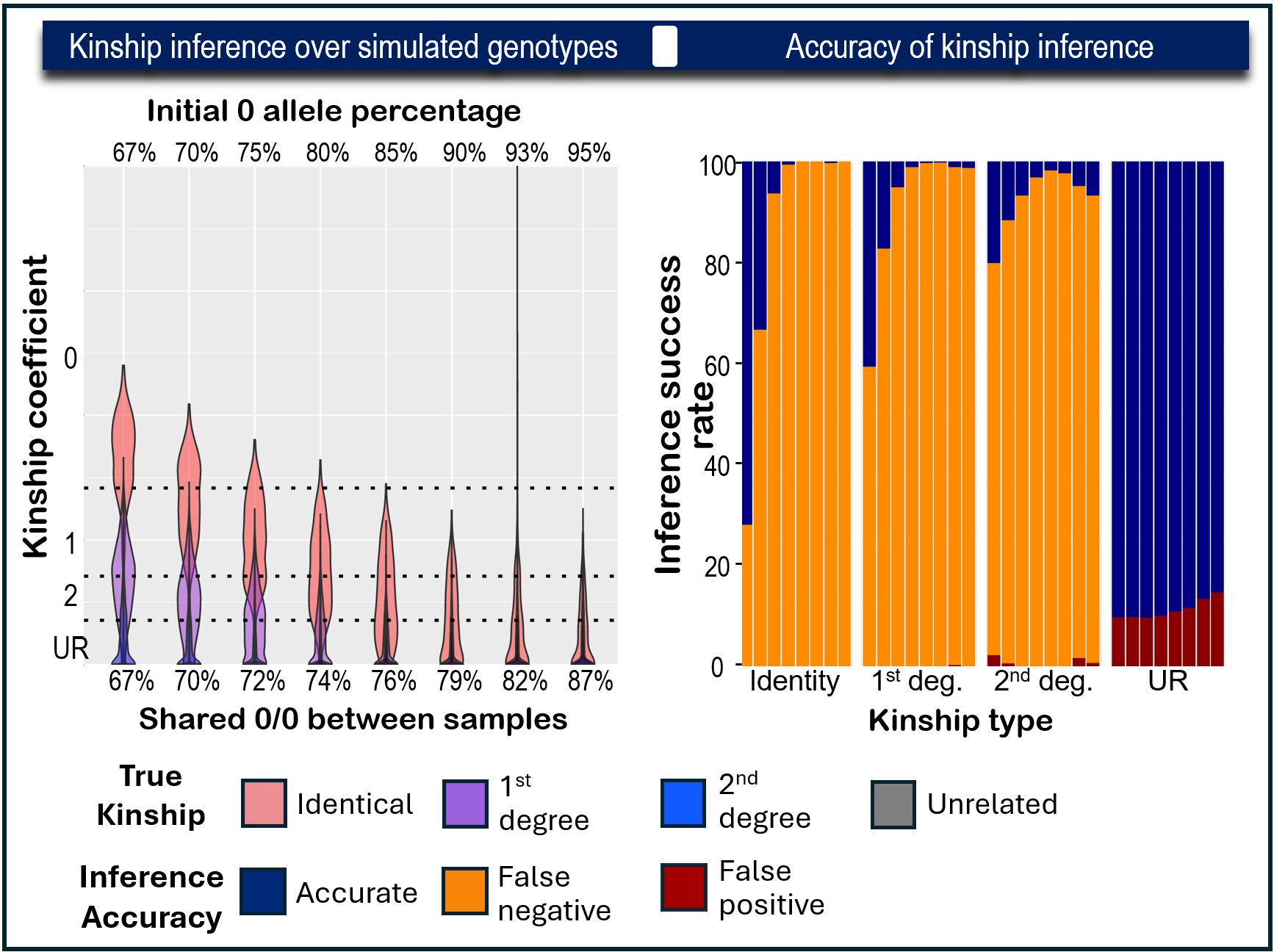
**Supplementary Figure S6.** Kinship coefficient and inference accuracy obtained by running READ over 100 independently simulated families over which reference bias was simulated at varying rates. **Left**: Distribution of inferred kinship coefficient values. On top, the proportions of 0 alleles across all genotypes and in the bottom - the average proportions of shared 0/0 alleles across all pairs. **Right**: Accuracy of kinship inference. Four panels correspond to the four possible kinship types (identical, first-degree, second-degree and unrelated; X-axis). Colored bars indicate the proportions of accurate, false negative and false positive inferences across the different proportions given in the left panel.

#### **Chapter 4.** A simple test to identify signs of mixture of individuals in a single sample

##### Description of methodology

We developed a straightforward test to assess whether a sedimentary sample contains genetic material from more than one individual. Intuitively, assuming a population with uniform heterozygosity level, a sample containing DNA from multiple individuals will show higher than expected heterozygosity levels. To assess whether heterozygosity could serve as a diagnostic signal for individual mixture, we examined the frequency of heterozygous loci, defined here as the ratio of heterozygous (0/1) to homozygous alternative (1/1) genotypes, where 0 is the reference allele and 1 is the alternative allele. To test this approach, we used *Sim1000G* to simulate 22 unrelated individuals and 10 full siblings from the Chagyrskaya population, as detailed in the Methods (section “Simulating a Neandertal family”). Modern human contamination was simulated using *ArchSim*, as described in the Methods (section “Simulating sedimentary ancient DNA data”).

Next, we created composite genotypes from by randomly selecting alleles from an increasing number of the simulated individuals and measured the resulting heterozygosity. To approximate biologically realistic mixture within a sample, the composite genotypes were generated by pooling alleles from the required number of individuals at each locus according to predefined proportions (e.g., a 90:10 mixture contributed nine alleles from one individual, chosen randomly, and a single allele from the other and an equal proportions mixture contributed one allele from each of the individuals mixed together), followed by a random sampling of two alleles from this pool to construct the composite genotype.

We checked the following scenarios:

1. The mixture of genotypes of up to eight individuals, which were randomly selected from 22 unrelated individuals simulated using the genotypes of the Chagyrskaya population. Mean simulated reads depth was 5.
2. The same as 1, but with the addition of simulated 10% of modern human contamination to each mixture.
3. The mixture of up to eight full siblings which were randomly selected from 10 full siblings simulated using the Chagyrskaya genotypes, with a mean read depth of 5.
4. The mixture of genotypes in chromosome 22 of up to eight individuals, which were randomly selected from the Finn population in the 1000 Genomes project. This population was chosen due to its low variability relative to other human populations in the 1000 Genome project phase III^28^.

In each scenario, each individual’s genotypes were mixed in equal proportions. In addition, two individuals were mixed with the proportions of 90% - 10% and 75% - 25%. For each number of mixed individuals, and each mixture proportion, these scenarios were simulated 100 times. We opted to assess the results only when the number of SNPs was at least 1,000.

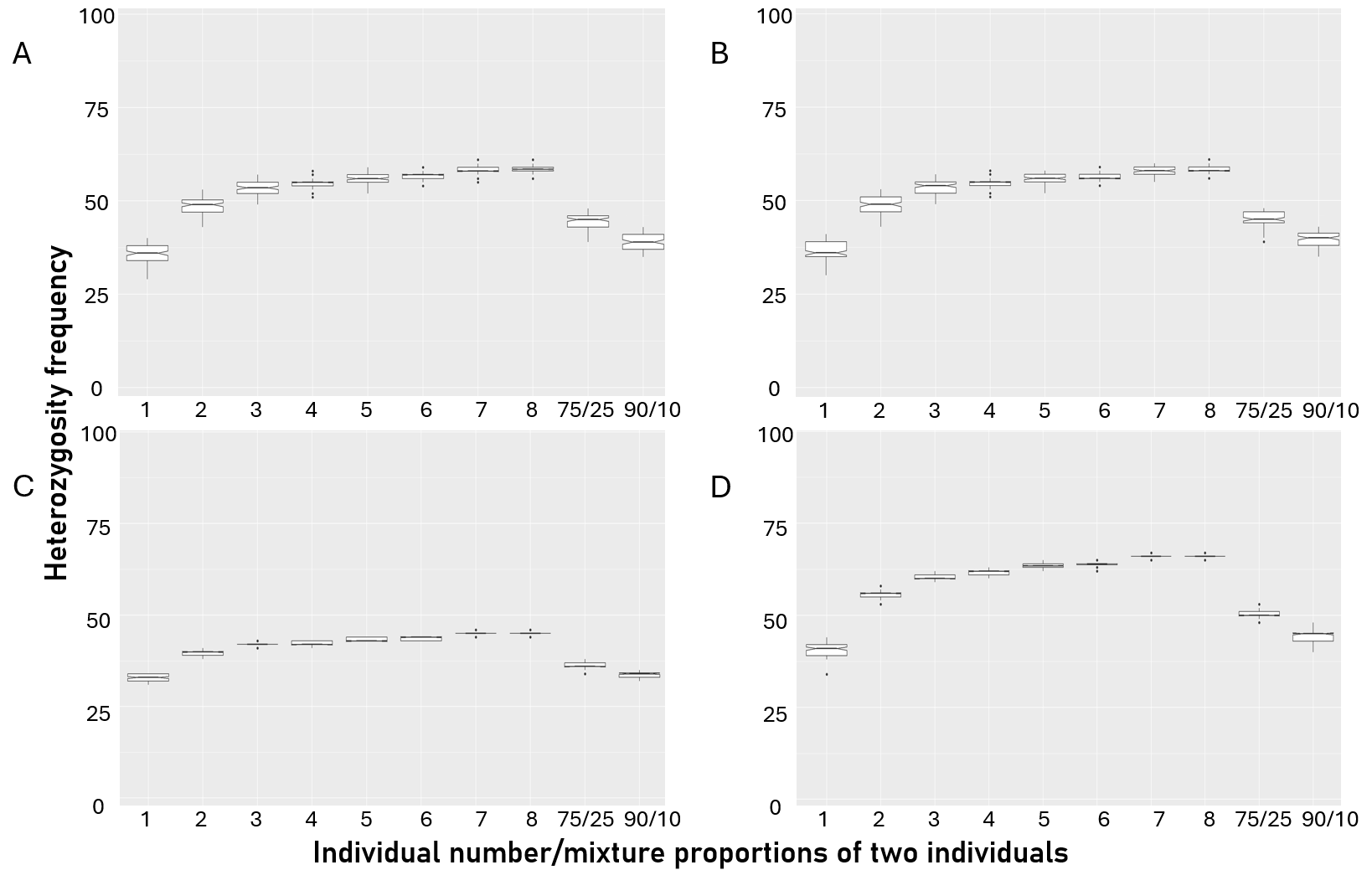
**Supplementary Figure S7**. Heterozygosity frequency measured in genotypes of varying number of individuals mixed in equal proportions, or for two individuals mixed in inequal proportions. **A**. Individuals were randomly selected from 22 unrelated individuals which were simulated using the genotypes inferred for the Chagyrskaya population. **B**. Same as A with added 10% of modern human contamination. **C**. Individuals were randomly selected from full siblings which were simulated using the genotypes inferred for the Chagyrskaya population. **D**. Individuals were randomly selected from the Finn population in the 1000 Genome project and only the genotypes on chromosome 22 were used.

##### Interpretations and uses

We observed a clear increase in heterozygosity as the number of individuals contributing to a sample increased, as expected (Supplemtary Figure S7). The most pronounced shift occurred between samples derived from a single individual and those representing a mixture of two individuals. In nearly all scenarios, heterozygosity levels from single-individual samples were non-overlapping with those of two-individual mixtures. Only when two individuals were combined in highly skewed proportions (e.g., 90% and 10%), heterozygosity levels partially overlapped those of a single individual.

These results indicate that heterozygosity is a simple and effective metric for distinguishing between single- and multi-individual samples, even in the presence of modern human contamination or when the individuals are closely related. However, this approach is robust only under two key conditions: (1) the comparison is performed within a single population that lacks significant substructuring, and (2) the baseline heterozygosity level of an individual from that population, under comparable preservation and sequencing conditions, is either known or can be reliably estimated.

We suggest that this heterozygosity frequency measure can be used to support or complement other methods intended for the same purpose, such as the reconstruction of mtDNA consensus sequence or comparative analysis of sex chromosomes mapped sequences (e.g., Slon et al., 2017; Zavala et al., 2021; Vernot et al., 2021)^1,29,30^. Indeed, heterozygosity levels can potentially detect the presence of multiple individuals even in cases where other approaches may not be able to do so; for example, when different individuals share the same mtDNA.

##### Application to the sedimentary samples from Galerìa de las Estatuas

To enable a preliminary assessment of whether the Estatuas sedimentary samples are more consistent with deriving from a single individual or from a mixture of individuals, we conducted a targeted simulation designed to approximate the characteristics of that dataset. Specifically, we repeated the simulation described as ‘scenario 1’ above (Supplementary Figure S7A) but limiting the analysis to SNPs that are present in the Estatuas dataset. This constraint resulted in only 77 usable SNPs, far below our set threshold of at least 1,000 SNPs for this analysis. As such, the results, shown in Supplementary Figure S8, should be interpreted with caution, as the limited number of SNPs reduces the robustness of the comparison.

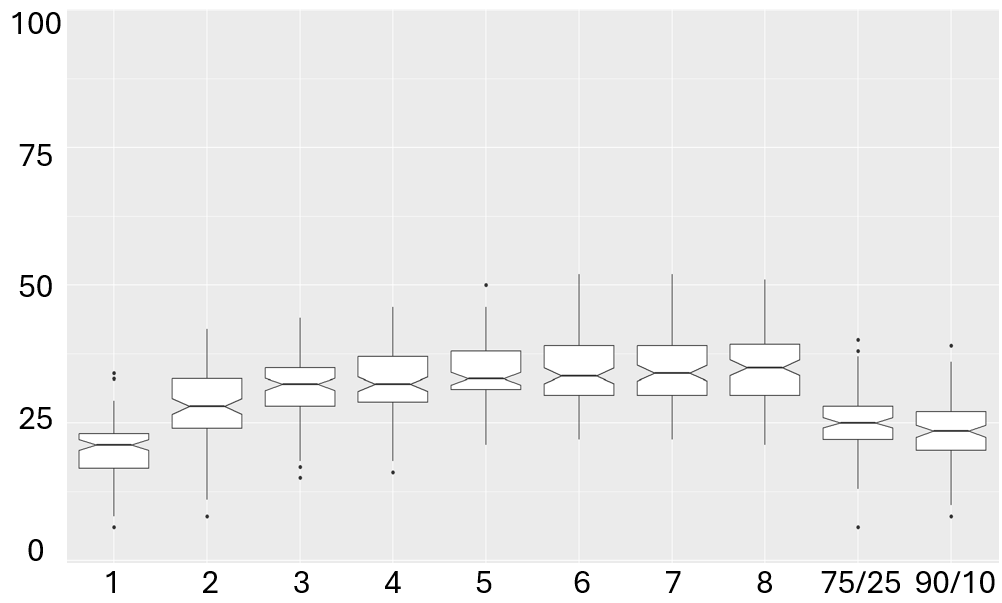
**Supplementary Figure S8**. Heterozygosity frequency measured in genotypes of varying number of individuals mixed together in equal proportions, or for two individuals in inequal proportions. The genotypes used in this illustration are only at SNPs present in the Estatuas Neandertal population.

#### **Chapter 5.** Samples from Galería de las Estatuas

##### Processing only putatively deaminated fragments

We downloaded the unmapped bam files of the nuclear-DNA enriched sedimentary samples that were collected in the Northern Spanish site of the Galería de las Estatuas and described in Vernot *et al* (2021)^1^. From each bam file, sequences shorter than 35bp were filtered out a priori using *samtools* to save computation time, and then mapped to the human reference genome (hg19 with ‘decoy’ sequences) using *BWA*^31^ with ‘ancient’ parameters^32^. Unmapped sequences and those with a mapping quality lower than 25 were filtered out using *samtools*. All bam files of the same sub-sample (identified by their library name) were merged using the *samtools* merge option, resulting in 105 samples. Duplicated sequences were removed based on sequence content. Reads that did not show signs of deamination were also removed (based on the presence of C-T substititions [G to A for fragments sequenced in reverse orientation] to the reference genome at the first three or last three bases). These putatively deaminated bases were masked. The samples were genotyped using the *HaplotypeCaller*, *CombineGVCFs* and *GenotypeGVCFs* protocols with the default parameters of the GATK pipeline^18^. This was done for all 105 samples together and for each sample separately, so heterozygosity level could be studied. The unfiltered combined genotype file contained 15,324 SNPs, while the separate genotype files contained between 1 and 2,243 SNPs. Each separate genotype file was filtered for a minimal read depth of 6 in the binominal test for composite samples leaving between 0 and 282 SNPs. Full processing results are in Supplementary Table S3.

**Supplementary Table S3.** Processing details of 105 sub-samples resulting from the merging of unmapped bam files downloaded from <https://www.ebi.ac.uk/ena/browser/view/PRJEB42656> The number of high-quality SNPs remaining after all processing and filtering steps is shown under ‘SPNs filtered’,

| **library** | **sample** | **sub-sample** | **Pit** | **Layer** | **Reads retained after bam processing** | **deaminated unique reads** | **SNPs** | **SNPs filtered** |
| --- | --- | --- | --- | --- | --- | --- | --- | --- |
| **A11419** | GE-I-A4 | GE-I-A4c | I | 4 | 238,349 | 28,687 | 662 | 50 |
| **A11421** | GE-I-A4 | GE-I-A4e | I | 4 | 240,516 | 29,697 | 739 | 31 |
| **A11422** | GE-I-A4 | GE-I-A4f | I | 4 | 236,801 | 24,383 | 536 | 26 |
| **A11423** | GE-I-A4 | GE-I-A4g | I | 4 | 175,985 | 25,090 | 746 | 78 |
| **A11424** | GE-I-A4 | GE-I-A4h | I | 4 | 203,044 | 21,477 | 441 | 29 |
| **A11426** | GE-I-A4 | GE-I-A4j | I | 4 | 30,637 | 5,278 | 79 | 5 |
| **A15864** | GE-I-A4 | GE-I-A4g | I | 4 | 89,764 | 11,832 | 208 | 16 |
| **A16042** | GE-I-B07 | GE-I-B07a | I | 2 | 27,708 | 4,253 | 107 | 5 |
| **A16043** | GE-I-B08 | GE-I-B08a | I | 2 | 24,919 | 3,879 | 67 | 1 |
| **A16044** | GE-I-B09 | GE-I-B09a | I | 2 | 254,840 | 30,516 | 1,224 | 223 |
| **A16045** | GE-I-B10 | GE-I-B10a | I | 3 | 615,296 | 65,217 | 2,243 | 217 |
| **A16046** | GE-I-B11 | GE-I-B11a | I | 3 | 307,972 | 32,530 | 1,143 | 162 |
| **A16060** | GE-I-B19 | GE-I-B19a | I | 3 | 205,534 | 28,862 | 747 | 50 |
| **A16070** | GE-I-B29 | GE-I-B29a | I | 2 | 43,697 | 6,739 | 221 | 15 |
| **A16071** | GE-I-B30 | GE-I-B30a | I | 3 | 472,855 | 34,477 | 1,043 | 115 |
| **A16072** | GE-I-B31 | GE-I-B31a | I | 3 | 42,853 | 6,900 | 222 | 17 |
| **A16073** | GE-I-B32 | GE-I-B32a | I | 3.5 | 557,572 | 52,058 | 1,828 | 282 |
| **A16074** | GE-I-B33 | GE-I-B33a | I | 4 | 20,780 | 2,633 | 32 | 2 |
| **A16075** | GE-I-B34 | GE-I-B34a | I | 3.5 | 33,204 | 4,416 | 60 | 2 |
| **A16084** | GE-I-B37 | GE-I-B37a | I | 4 | 18,721 | 2,116 | 25 | 2 |
| **A16085** | GE-I-B38 | GE-I-B38b | I | 5 | 60,282 | 2,964 | 39 | 3 |
| **A16089** | GE-I-B42 | GE-I-B42b | I | 4 | 206,848 | 15,449 | 403 | 43 |
| **A16091** | GE-I-B44 | GE-I-B44a | I | 3.5 | 41,186 | 4,254 | 60 | 4 |
| **A16092** | GE-I-B45 | GE-I-B45b | I | 4 | 63,823 | 4,679 | 79 | 9 |
| **A16093** | GE-I-B46 | GE-I-B46a | I | 4 | 47,625 | 3,084 | 50 | 0 |
| **A16094** | GE-I-B47 | GE-I-B47b | I | 4 | 19,836 | 2,003 | 7 | 0 |
| **A16095** | GE-I-B48 | GE-I-B48a | I | 5 | 42,162 | 2,267 | 23 | 2 |
| **A16096** | GE-II-B100 | GE-II-B100a | II | 2.5 | 71975 | 474 | 1 | 0 |
| **A16097** | GE-II-B101 | GE-II-B101a | II | 2 | 112,365 | 1,658 | 18 | 2 |
| **A16098** | GE-II-B102 | GE-II-B102a | II | 2 | 106,419 | 1,287 | 11 | 2 |
| **A16100** | GE-II-B104 | GE-II-B104a | II | 2 | 46,681 | 5,175 | 107 | 7 |
| **A16101** | GE-II-B105 | GE-II-B105a | II | 2 | 57,241 | 8,488 | 265 | 26 |
| **A16109** | GE-II-B107 | GE-II-B107a | II | 2 | 118,251 | 2,736 | 37 | 2 |
| **A16110** | GE-II-B108 | GE-II-B108a | II | 2 | 314,208 | 33,398 | 1,213 | 205 |
| **A16111** | GE-II-B109 | GE-II-B109a | II | 2 | 620,892 | 56,790 | 1,750 | 244 |
| **A16112** | GE-II-B110 | GE-II-B110a | II | 2 | 159,516 | 28,784 | 936 | 81 |
| **A20280** | GE-I-A4 | GE-I-A4k | I | 4 | 130,590 | 18,433 | 608 | 56 |
| **A20281** | GE-I-A4 | GE-I-A4l | I | 4 | 185,415 | 23,364 | 757 | 87 |
| **A20282** | GE-I-A4 | GE-I-A4n | I | 4 | 123,485 | 16,815 | 438 | 48 |
| **A20285** | GE-I-A4 | GE-I-A4o | I | 4 | 103,880 | 12,421 | 318 | 23 |
| **A20286** | GE-I-A4 | GE-I-A4m | I | 4 | 134,036 | 18,160 | 581 | 54 |
| **A20287** | GE-I-B09 | GE-I-B09a | I | 2 | 95,110 | 14,506 | 387 | 20 |
| **A20293** | GE-I-B42 | GE-I-B42a | I | 4 | 111,923 | 8,359 | 158 | 11 |
| **A20301** | GE-I-B47 | GE-I-B47a | I | 4 | 93,496 | 4,967 | 50 | 6 |
| **A20322** | GE-I-B08 | GE-I-B08b | I | 2 | 189,474 | 21,445 | 605 | 37 |
| **A20323** | GE-I-B08 | GE-I-B08c | I | 2 | 88,878 | 11,138 | 279 | 19 |
| **A20324** | GE-I-B08 | GE-I-B08d | I | 2 | 92,326 | 12,051 | 356 | 24 |
| **A20332** | GE-I-B18 | GE-I-B18c | I | 2 | 100,239 | 12,839 | 283 | 15 |
| **A20347** | GE-I-B32 | GE-I-B32b | I | 3.5 | 109,084 | 15,269 | 449 | 53 |
| **A20348** | GE-I-B32 | GE-I-B32c | I | 3.5 | 131,567 | 16,919 | 522 | 54 |
| **A20362** | GE-I-B37 | GE-I-B37b | I | 4 | 68,461 | 7,999 | 134 | 11 |
| **A20371** | GE-I-B38 | GE-I-B38f | I | 5 | 34,645 | 3,201 | 17 | 3 |
| **A20372** | GE-I-B39 | GE-I-B39b | I | 3.5 | 61,945 | 9,352 | 157 | 11 |
| **A20376** | GE-I-B40 | GE-I-B40d | I | 4 | 63,391 | 6,814 | 101 | 11 |
| **A20384** | GE-I-B42 | GE-I-B42e | I | 4 | 105,301 | 14,667 | 379 | 24 |
| **A20386** | GE-I-B43 | GE-I-B43b | I | 5 | 55,639 | 3,360 | 32 | 5 |
| **A20392** | GE-I-B45 | GE-I-B45c | I | 4 | 73,770 | 8,025 | 156 | 10 |
| **A20396** | GE-I-B45 | GE-I-B45d | I | 4 | 82,464 | 9,516 | 197 | 11 |
| **A20398** | GE-I-B45 | GE-I-B45f | I | 4 | 93,192 | 9,055 | 132 | 11 |
| **A20403** | GE-I-B47 | GE-I-B47d | I | 4 | 155,968 | 12,505 | 223 | 18 |
| **A24113** | GE-I-B07 | GE-I-B07f | I | 2 | 71,264 | 5,978 | 91 | 7 |
| **A24114** | GE-I-B07 | GE-I-B07g | I | 2 | 86,116 | 7,123 | 143 | 7 |
| **A24115** | GE-I-B07 | GE-I-B07h | I | 2 | 52,751 | 5,026 | 88 | 9 |
| **A24116** | GE-I-B08 | GE-I-B08f | I | 2 | 119,535 | 12,599 | 304 | 30 |
| **A24117** | GE-I-B08 | GE-I-B08g | I | 2 | 92,662 | 9,625 | 231 | 11 |
| **A24118** | GE-I-B08 | GE-I-B08h | I | 2 | 93,632 | 5,264 | 80 | 6 |
| **A24119** | GE-I-B18 | GE-I-B18f | I | 2 | 121,277 | 11,360 | 284 | 25 |
| **A24120** | GE-I-B18 | GE-I-B18g | I | 2 | 86,114 | 9,061 | 181 | 15 |
| **A24121** | GE-I-B18 | GE-I-B18h | I | 2 | 86,236 | 8,792 | 212 | 18 |
| **A24122** | GE-I-B29 | GE-I-B29f | I | 2 | 102,406 | 12,439 | 383 | 26 |
| **A24125** | GE-I-B29 | GE-I-B29g | I | 2 | 212,946 | 28,937 | 953 | 110 |
| **A24126** | GE-I-B29 | GE-I-B29h | I | 2 | 94,230 | 14,714 | 519 | 61 |
| **A24127** | GE-I-B33 | GE-I-B33f | I | 4 | 78,316 | 5,751 | 102 | 8 |
| **A24128** | GE-I-B33 | GE-I-B33g | I | 4 | 57,588 | 4,603 | 74 | 5 |
| **A24129** | GE-I-B33 | GE-I-B33h | I | 4 | 93,860 | 6,309 | 101 | 5 |
| **A24131** | GE-I-B37 | GE-I-B37f | I | 4 | 77,299 | 6,126 | 84 | 12 |
| **A24132** | GE-I-B37 | GE-I-B37g | I | 4 | 77,814 | 6,528 | 120 | 14 |
| **A24133** | GE-I-B37 | GE-I-B37h | I | 4 | 52,195 | 5,340 | 92 | 9 |
| **A24134** | GE-I-B38 | GE-I-B38g | I | 5 | 434,906 | 4,348 | 82 | 14 |
| **A24135** | GE-I-B38 | GE-I-B38h | I | 5 | 52,312 | 3,929 | 58 | 8 |
| **A24136** | GE-I-B38 | GE-I-B38i | I | 5 | 243,590 | 4,958 | 69 | 12 |
| **A24137** | GE-I-B40 | GE-I-B40f | I | 4 | 92,869 | 7,517 | 117 | 7 |
| **A24138** | GE-I-B40 | GE-I-B40g | I | 4 | 58,249 | 4,828 | 68 | 5 |
| **A24139** | GE-I-B40 | GE-I-B40h | I | 4 | 73,988 | 6,075 | 94 | 12 |
| **A24140** | GE-I-B42 | GE-I-B42g | I | 4 | 132,401 | 12,764 | 404 | 36 |
| **A24141** | GE-I-B42 | GE-I-B42h | I | 4 | 60,220 | 5,481 | 91 | 18 |
| **A24142** | GE-I-B42 | GE-I-B42i | I | 4 | 57,857 | 5,728 | 92 | 10 |
| **A24143** | GE-I-B43 | GE-I-B43f | I | 5 | 69,973 | 3,479 | 35 | 0 |
| **A24144** | GE-I-B43 | GE-I-B43g | I | 5 | 102,281 | 5,023 | 58 | 8 |
| **A24145** | GE-I-B43 | GE-I-B43h | I | 5 | 63,352 | 3,684 | 47 | 3 |
| **A24150** | GE-I-B45 | GE-I-B45g | I | 4 | 96,518 | 11,437 | 251 | 33 |
| **A24151** | GE-I-B45 | GE-I-B45h | I | 4 | 83,639 | 6,465 | 105 | 10 |
| **A24152** | GE-I-B45 | GE-I-B45i | I | 4 | 88,831 | 7,095 | 120 | 21 |
| **A24153** | GE-I-B47 | GE-I-B47g | I | 4 | 35,090 | 3,950 | 47 | 5 |
| **A24154** | GE-I-B47 | GE-I-B47h | I | 4 | 51,990 | 4,733 | 79 | 8 |
| **A24155** | GE-I-B47 | GE-I-B47i | I | 4 | 55,835 | 4,990 | 74 | 5 |
| **A24156** | GE-I-B48 | GE-I-B48f | I | 5 | 51,250 | 3,401 | 28 | 2 |
| **A24157** | GE-I-B48 | GE-I-B48g | I | 5 | 62,969 | 3,433 | 32 | 0 |
| **A24158** | GE-I-B48 | GE-I-B48h | I | 5 | 40,712 | 3,382 | 27 | 4 |
| **A24518** | GE-I-A4 | GE-I-A4l | I | 4 | 250,316 | 25,330 | 809 | 79 |
| **A24519** | GE-I-A4 | GE-I-A4l | I | 4 | 212,879 | 21,589 | 755 | 57 |
| **A24756** | GE-I-A4 | GE-I-A4l | I | 4 | 228,718 | 25,519 | 871 | 73 |
| **A28210** | GE-I-B43 | GE-I-B43i | I | 5 | 27,189 | 509 | 9 | 0 |
| **A28219** | GE-I-B43 | GE-I-B43j | I | 5 | 17048 | 450 | 7 | 0 |
| **A28238** | GE-I-B43 | GE-I-B43k | I | 5 | 28,967 | 1,635 | 69 | 2 |

##### Binomial test to determine whether samples contain DNA from a single or multiple individuals

A one-tailed binomial test was carried out over 105 Galerìa de las Estatuas samples (using putatively deaminated sequences only) with parameters (*N, p_0_*). *N* is the total number of sites carrying the alternative allele genotyped as either heterozygous (0/1) or homozygous alternative (1/1) and *S* = the number of heterozygous sites (0/1). The observed heterozygosity frequency was calculated as *S*/*N*. *p_0_* is baseline heterozygosity expectation for an individual in the population. Three values are suggested for *p_0_* : 0.21 - derived from the simulations of heterozygosity using genotypes inferred for the Chagyrskaya population and shared by those inferred for the Estatuas population, 0.36 and 0.4 (derived from the simulations of heterozygosity using genotypes inferred for the Chagyrskaya population - median and maximal value respectively). See Supplementary Figures S7 and S8.

**Supplementary Table S4**. Binomial test results for 105 Galerìa de las Estatuas sedimentary samples under p₀ = 0.21, 0.36, and 0.40. The null model, for which a sample derives from a single individual, is rejected when the observed number of heterozygous SNPs is higher than expected for one individual under the corresponding *p₀*. One, two, and three asterisks indicate rejection under *p₀* = 0.21, 0.36, and 0.40, respectively. Colored entries mark samples whose consensus mtDNA sequences were classified by Vernot *et al*. (2021) as likely originating from either a single hominin (green) or multiple hominins (blue)^1^.

| **library** | **SNPs** | **SNPs filtered** | **heterozygote SNPs** | **p-value (p0=0.21)** | **p-value (p0=0.36)** | **p-value (p0=0.4)** |
| --- | --- | --- | --- | --- | --- | --- |
| **A11419**** | 662 | 50 | 25 | 5.28E-06 | 0.029483897 | 0.09780736 |
| **A11421*** | 739 | 31 | 16 | 0.00016202 | 0.054371766 | 0.12838173 |
| **A11422*** | 536 | 26 | 13 | 0.00098298 | 0.101303384 | 0.19934779 |
| **A11423***** | 746 | 78 | 40 | 3.49E-09 | 0.004078144 | 0.02847972 |
| **A11424***** | 441 | 29 | 17 | 1.11E-05 | 0.010825734 | 0.03288434 |
| **A11426** | 79 | 5 | 2 | 0.28331851 | 0.59063593 | 0.66304 |
| **A15864*** | 208 | 16 | 7 | 0.03420217 | 0.342815161 | 0.47282589 |
| **A16042** | 107 | 5 | 0 | 1 | 1 | 1 |
| **A16043** | 67 | 1 | 0 | 1 | 1 | 1 |
| **A16044*** | 1,224 | 223 | 82 | 4.85E-08 | 0.429962375 | 0.8538509 |
| **A16045*** | 2,243 | 217 | 71 | 3.93E-05 | 0.859679955 | 0.9887568 |
| **A16046*** | 1,143 | 162 | 49 | 0.00358393 | 0.947535679 | 0.9960384 |
| **A16060*** | 747 | 50 | 24 | 1.96E-05 | 0.054412017 | 0.15616833 |
| **A16070*** | 221 | 15 | 8 | 0.00581491 | 0.13022007 | 0.21310318 |
| **A16071**** | 1,043 | 115 | 56 | 4.11E-11 | 0.003516678 | 0.03615453 |
| **A16072*** | 222 | 17 | 7 | 0.04785838 | 0.415165623 | 0.55215937 |
| **A16073**** | 1,828 | 282 | 117 | 6.28E-15 | 0.032450405 | 0.32531781 |
| **A16074** | 32 | 2 | 0 | 1 | 1 | 1 |
| **A16075** | 60 | 2 | 0 | 1 | 1 | 1 |
| **A16084** | 25 | 2 | 0 | 1 | 1 | 1 |
| **A16085** | 39 | 3 | 1 | 0.506961 | 0.737856 | 0.784 |
| **A16089***** | 403 | 43 | 25 | 1.21E-07 | 0.002535573 | 0.01228048 |
| **A16091** | 60 | 4 | 1 | 0.61049919 | 0.83222784 | 0.8704 |
| **A16092** | 79 | 9 | 0 | 1 | 1 | 1 |
| **A16093** | 50 | 0 | 0 | 1 | 1 | 1 |
| **A16094** | 7 | 0 | 0 | 1 | 1 | 1 |
| **A16095** | 23 | 2 | 0 | 1 | 1 | 1 |
| **A16096** | 1 | 0 | 0 | 1 | 1 | 1 |
| **A16097** | 18 | 2 | 0 | 1 | 1 | 1 |
| **A16098** | 11 | 2 | 0 | 1 | 1 | 1 |
| **A16100** | 107 | 7 | 0 | 1 | 1 | 1 |
| **A16101** | 265 | 26 | 7 | 0.29632868 | 0.880567559 | 0.94411596 |
| **A16109** | 37 | 2 | 0 | 1 | 1 | 1 |
| **A16110*** | 1,213 | 205 | 83 | 2.03E-10 | 0.103424287 | 0.46971291 |
| **A16111*** | 1,750 | 244 | 100 | 1.31E-12 | 0.060896878 | 0.40040523 |
| **A16112*** | 936 | 81 | 26 | 0.01304078 | 0.800646246 | 0.94282249 |
| **A20280***** | 608 | 56 | 35 | 2.13E-11 | 4.96E-05 | 0.00055652 |
| **A20281***** | 757 | 87 | 52 | 4.55E-15 | 5.52E-06 | 0.00015365 |
| **A20282***** | 438 | 48 | 28 | 1.93E-08 | 0.001333528 | 0.00779379 |
| **A20285** | 318 | 23 | 6 | 0.34967455 | 0.888824459 | 0.94603143 |
| **A20286**** | 581 | 54 | 27 | 2.24E-06 | 0.024266856 | 0.08766781 |
| **A20287** | 387 | 20 | 7 | 0.10710115 | 0.619685286 | 0.74998933 |
| **A20293** | 158 | 11 | 2 | 0.70648344 | 0.946965611 | 0.96976691 |
| **A20301** | 50 | 6 | 3 | 0.11154865 | 0.37320321 | 0.45568 |
| **A20322** | 605 | 37 | 12 | 0.07103344 | 0.729865871 | 0.86665092 |
| **A20323** | 279 | 19 | 5 | 0.368057 | 0.86992428 | 0.93038629 |
| **A20324*** | 356 | 24 | 13 | 0.00036115 | 0.052709839 | 0.11426508 |
| **A20332*** | 283 | 15 | 9 | 0.00114087 | 0.050378628 | 0.09504741 |
| **A20347*** | 449 | 53 | 21 | 0.00156575 | 0.338325055 | 0.57412199 |
| **A20348**** | 522 | 54 | 27 | 2.24E-06 | 0.024266856 | 0.08766781 |
| **A20362** | 134 | 11 | 3 | 0.4157834 | 0.818558978 | 0.88108319 |
| **A20371** | 17 | 3 | 0 | 1 | 1 | 1 |
| **A20372** | 157 | 11 | 3 | 0.4157834 | 0.818558978 | 0.88108319 |
| **A20376** | 101 | 11 | 5 | 0.06071106 | 0.358100819 | 0.4672258 |
| **A20384*** | 379 | 24 | 11 | 0.00561819 | 0.212516198 | 0.34976187 |
| **A20386** | 32 | 5 | 3 | 0.06588831 | 0.250897306 | 0.31744 |
| **A20392*** | 156 | 10 | 5 | 0.03986239 | 0.270841536 | 0.36689674 |
| **A20396*** | 197 | 11 | 6 | 0.01484399 | 0.166130396 | 0.24650187 |
| **A20398** | 132 | 11 | 3 | 0.4157834 | 0.818558978 | 0.88108319 |
| **A20403** | 223 | 18 | 3 | 0.7616075 | 0.980679737 | 0.99177364 |
| **A24113** | 91 | 7 | 1 | 0.80796091 | 0.956019535 | 0.9720064 |
| **A24114** | 143 | 7 | 3 | 0.16565616 | 0.490616879 | 0.580096 |
| **A24115** | 88 | 9 | 3 | 0.28853358 | 0.685592451 | 0.76821299 |
| **A24116*** | 304 | 30 | 12 | 0.0139938 | 0.388880872 | 0.5689095 |
| **A24117** | 231 | 11 | 5 | 0.06071106 | 0.358100819 | 0.4672258 |
| **A24118** | 80 | 6 | 3 | 0.11154865 | 0.37320321 | 0.45568 |
| **A24119***** | 284 | 25 | 15 | 2.51E-05 | 0.012404725 | 0.03439152 |
| **A24120*** | 181 | 15 | 8 | 0.00581491 | 0.13022007 | 0.21310318 |
| **A24121** | 212 | 18 | 6 | 0.15861463 | 0.677556632 | 0.79124163 |
| **A24122**** | 383 | 26 | 14 | 0.00023298 | 0.047682921 | 0.10818775 |
| **A24125*** | 953 | 110 | 37 | 0.00143173 | 0.728962575 | 0.9290325 |
| **A24126*** | 519 | 61 | 25 | 0.00032029 | 0.247034969 | 0.4860921 |
| **A24127** | 102 | 8 | 2 | 0.52566345 | 0.845188763 | 0.89362432 |
| **A24128** | 74 | 5 | 3 | 0.06588831 | 0.250897306 | 0.31744 |
| **A24129** | 101 | 5 | 1 | 0.69229436 | 0.892625818 | 0.92224 |
| **A24131** | 84 | 12 | 5 | 0.08659319 | 0.445858727 | 0.56182178 |
| **A24132** | 120 | 14 | 5 | 0.15226177 | 0.607988537 | 0.72074301 |
| **A24133** | 92 | 9 | 3 | 0.28853358 | 0.685592451 | 0.76821299 |
| **A24134*** | 82 | 14 | 7 | 0.01519267 | 0.205921586 | 0.3075478 |
| **A24135** | 58 | 8 | 3 | 0.22549906 | 0.595819526 | 0.68460544 |
| **A24136** | 69 | 12 | 2 | 0.75241405 | 0.96340166 | 0.98040896 |
| **A24137** | 117 | 7 | 3 | 0.16565616 | 0.490616879 | 0.580096 |
| **A24138** | 68 | 5 | 2 | 0.28331851 | 0.59063593 | 0.66304 |
| **A24139*** | 94 | 12 | 6 | 0.02447607 | 0.235239748 | 0.33479144 |
| **A24140*** | 404 | 36 | 15 | 0.00402537 | 0.292691933 | 0.4818917 |
| **A24141** | 91 | 18 | 6 | 0.15861463 | 0.677556632 | 0.79124163 |
| **A24142** | 92 | 10 | 4 | 0.13914176 | 0.513228434 | 0.6177194 |
| **A24143** | 35 | 0 | 0 | 1 | 1 | 1 |
| **A24144** | 58 | 8 | 3 | 0.22549906 | 0.595819526 | 0.68460544 |
| **A24145** | 47 | 3 | 2 | 0.113778 | 0.295488 | 0.352 |
| **A24150***** | 251 | 33 | 23 | 2.53E-09 | 8.60E-05 | 0.00053596 |
| **A24151** | 105 | 10 | 2 | 0.65362889 | 0.92361895 | 0.9536426 |
| **A24152*** | 120 | 21 | 11 | 0.00148485 | 0.092556027 | 0.17437787 |
| **A24153** | 47 | 5 | 0 | 1 | 1 | 1 |
| **A24154** | 79 | 8 | 2 | 0.52566345 | 0.845188763 | 0.89362432 |
| **A24155** | 74 | 5 | 1 | 0.69229436 | 0.892625818 | 0.92224 |
| **A24156** | 28 | 2 | 0 | 1 | 1 | 1 |
| **A24157** | 32 | 0 | 0 | 1 | 1 | 1 |
| **A24158** | 27 | 4 | 2 | 0.19634643 | 0.45474048 | 0.5248 |
| **A24518*** | 809 | 79 | 33 | 2.42E-05 | 0.170362198 | 0.41533509 |
| **A24519** | 755 | 57 | 23 | 0.00071025 | 0.289478023 | 0.52868163 |
| **A24756*** | 871 | 73 | 30 | 7.99E-05 | 0.214976149 | 0.46829413 |
| **A28210** | 9 | 0 | 0 | 1 | 1 | 1 |
| **A28219** | 7 | 0 | 0 | 1 | 1 | 1 |
| **A28238** | 69 | 2 | 0 | 1 | 1 | 1 |

##### Kinship Analysis

**Supplementary Table S5.** Kinship results inferred by *READ* analysis over 27 samples from Galerìa de las Estatuas. Only pairs sharing 800 SNPs and more are presented.

| **ID1** | **ID2** | **Rel** | **Zup** | **Zdown** | **P0 mean** | **Non normalized P0** | **Non normalized P0 serr** | **Overlap NSNPs** | **Kinship Coefficient** |
| --- | --- | --- | --- | --- | --- | --- | --- | --- | --- |
| A16045 | A16112 | Second Degree | 0.247 | 1.136 | 44 | 0.1902 | 0.0145 | 878 | 0.1105 |
| A11421 | A24125 | Unrelated | NA | 2.286 | 1.0836 | 0.2317 | 0.0166 | 820 | -0.0836 |
| A11419 | A24125 | Unrelated | NA | 2.502 | 1.0888 | 0.2328 | 0.0156 | 859 | -0.0888 |
| A16071 | A16112 | Unrelated | NA | 2.644 | 1.0980 | 0.2348 | 0.0155 | 920 | -0.0980 |
| A16110 | A16112 | Unrelated | NA | 2.522 | 1.0994 | 0.2351 | 0.0164 | 889 | -0.0994 |
| A11419 | A11421 | Unrelated | NA | 3.558 | 1.1704 | 0.2503 | 0.0159 | 915 | -0.1704 |
| A16071 | A20322 | Unrelated | NA | 4.023 | 1.2148 | 0.2598 | 0.0164 | 870 | -0.2148 |
| A16045 | A16110 | Unrelated | NA | 4.724 | 1.2193 | 0.2607 | 0.0142 | 1166 | -0.2193 |
| A16071 | A24518 | Unrelated | NA | 3.993 | 1.2312 | 0.2633 | 0.0174 | 828 | -0.2312 |
| A16110 | A24125 | Unrelated | NA | 4.411 | 1.2380 | 0.2647 | 0.0161 | 933 | -0.2380 |
| A11419 | A11422 | Unrelated | NA | 4.243 | 1.2426 | 0.2657 | 0.0170 | 843 | -0.2426 |
| A16071 | A24125 | Unrelated | NA | 4.861 | 1.2645 | 0.2704 | 0.0158 | 1017 | -0.2645 |
| A16045 | A16071 | Unrelated | NA | 6.009 | 1.3253 | 0.2834 | 0.0149 | 1175 | -0.3253 |
| A11421 | A11422 | Unrelated | NA | 5.186 | 1.3287 | 0.2841 | 0.0174 | 806 | -0.3287 |
| A16071 | A24756 | Unrelated | NA | 5.520 | 1.3748 | 0.2940 | 0.0181 | 813 | -0.3748 |
| A11419 | A16110 | Unrelated | NA | 6.583 | 1.4109 | 0.3017 | 0.0164 | 1001 | -0.4109 |
| A11422 | A16110 | Unrelated | NA | 7.611 | 1.5233 | 0.3257 | 0.0173 | 921 | -0.5233 |
| A11421 | A16110 | Unrelated | NA | 8.090 | 1.5285 | 0.3268 | 0.0164 | 976 | -0.5285 |
| A11421 | A16071 | Unrelated | NA | 8.207 | 1.5334 | 0.3279 | 0.0163 | 1104 | -0.5334 |
| A11419 | A16071 | Unrelated | NA | 8.377 | 1.5368 | 0.3286 | 0.0161 | 1135 | -0.5368 |
| A16045 | A16060 | Unrelated | NA | 8.040 | 1.6054 | 0.3433 | 0.0186 | 836 | -0.6054 |
| A11422 | A16071 | Unrelated | NA | 9.604 | 1.6653 | 0.3561 | 0.0169 | 1025 | -0.6653 |
| A16071 | A16110 | Unrelated | NA | 11.294 | 1.6674 | 0.3565 | 0.0144 | 1551 | -0.6674 |
